# Substitution rate variation, not hidden paralogy, drives false hybridization signal in phylogenetic network inference

**DOI:** 10.64898/2026.05.11.723986

**Authors:** Bing Li, Cécile Ané

## Abstract

Phylogenetic network inference methods are increasingly used to detect hybridization and gene flow from genomic data, but their robustness to common sources of model violation remains poorly characterized. We conducted a simulation study to evaluate the effects of hidden paralogy and substitution rate variation on three commonly used network inference methods: find_graphs from ADMIXTOOLS 2, SNaQ and PhyNEST. Using eight-taxon species trees calibrated from empirical reptile and plant phylogenies, we simulated data under various levels of hidden paralogy (from none to strong) and three levels of rate variation (none, gene-specific, and lineage-specific). We found that hidden paralogy had limited impact on network inference under the conditions examined. In contrast, lineage-specific rates severely biased find_graphs, inflating worst *f* -statistic residuals well beyond the standard acceptance threshold. Under lineage-specific rates, PhyNEST had high type I error rates and was less likely to recover the true species topologies. SNaQ mostly selected the correct tree model, even under lineage-specific rates. Its networks inferred generally displayed the true tree, although recovery of the reptile tree declined under lineage-specific rates. We also show that the standard worst residuals threshold of 3 for find_graphs produces inflated type I error even without rate variation, as did the calibrated thresholds for PhyNEST, which differed between species trees and ILS levels. We therefore recommend empirical calibration of the model-selection threshold within each study system, and recommend caution using find_graphs and PhyNEST when rate variation across lineages is suspected.

## 1. Introduction

Traditional phylogenetic trees represent an evolutionary history as a bifurcating branching pattern, which reduces diversification to a series of divergence from common ancestors. Hybridization and introgression have been documented across a broad taxonomic range (Goulet et al., 2017; Mallet et al., 2016), from flowering plants such as oaks and poplars (Muir et al., 2000; Wang et al., 2020) to reptiles such as sea turtles and lizards (Barley et al., 2022; Pavón-Vázquez et al., 2021; Vilaça et al., 2021). Such reticulation processes generate complex patterns that cannot be adequately represented by simple bifurcating trees (Hibbins and Hahn, 2022). Phylogenetic networks and admixture graphs address this limitation, using extra edges to represent reticulation events (Huson and Scornavacca, 2011; Lipson, 2020). Consequently, phylogenetic network inference has become increasingly important in evolutionary biology and phylogenomics for detecting historical gene flow and reticulate evolution.

A wide variety of approaches have been developed to infer reticulate evolutionary histories, ranging from summary statistics derived from allele frequency or gene tree topologies to likelihood models explicitly reconstructing full phylogenetic networks (Hibbins and Hahn, 2022). Simpler tests based on summary statistics, such as the *D*-statistic (Green et al., 2010), *D*_3_-statistic (Hahn and Hibbins, 2019), and *f* -statistics (Patterson et al., 2012), measure asymmetries in shared derived alleles across taxa that are inconsistent with a strictly bifurcating species tree. These tests are computationally efficient approaches for detecting the presence of gene flow. More recently, many methods have been developed to explicitly model the multispecies coalescent under a species network. Full-likelihood methods include maximum-likelihood implementations, for example in PhyloNet (Yu et al., 2014) and Legofit (Rogers, 2019); as well as Bayesian methods for example in PhyloNet (e.g. Wen et al., 2018), BPP (Flouri et al., 2020) and AdmixtureBayes (Nielsen et al., 2023). More computationally efficient pseudolikelihood approaches include InferNetwork_MPL in PhyloNet (Yu and Nakhleh, 2015), SNaQ (Solís-Lemus and Ané, 2016), find_graphs (Maier et al., 2023) and PhyNEST (Kong et al., 2025). These methods differ in scope: some aim to infer the full network topology (e.g. SNaQ, find_graphs and PhyNEST), whereas others focus on estimating model parameters over a restricted set of candidate network topologies (e.g. BPP and Legofit).

In this study, we focus on three network estimation methods: find_graphs (implemented in the R package ADMIXTOOLS 2 v.2.0.10, Maier et al., 2023), SNaQ v.1.0.0 (a Julia package depending on PhyloNetworks v1.0, Solís-Lemus and Ané, 2016; Solís-Lemus et al., 2017), and PhyNEST v.0.1.12 (a Julia package, Kong et al., 2025). All three methods score candidate networks using summaries of data from small subsets of taxa, which explains their computational efficiency and wide adoption. find_graphs fits *f* -statistics, while SNaQ and PhyNEST combine likelihood contributions across four-taxon quartets. These methods account for incompletely lineage sorting, and jointly infer the network topology (including admixture direction) together with and edge parameters: edge lengths and gene flow proportions, or inheritance probabilities. They differ in their underlying data and model: find_graphs uses biallelic frequency data and a constant-rate model; SNaQ uses gene tree topologies, that need to be inferred from sequence data across predefined loci, and PhyNEST uses site pattern frequencies computed directly from the sequence alignment, without estimating gene trees. Together, the three methods span the main types of input used for network inference, which allows us to assess how sensitivity to model violations differs among input types.

In find_graphs, the pseudolikelihood score is based on *f* -statistics from genome-wide allele frequencies. It is optimized broadly over admixture graph topologies without restricting network complexity, allowing for networks of arbitrary level. However, identifiability of phylogenetic networks is generally guaranteed only for restricted classes, most notably level-1 networks, where reticulation events are sufficiently separated to induce non-overlapping cycles (Allman et al., 2024; Gross et al., 2021; Solís-Lemus and Ané, 2016). As network complexity increases, distinct topologies can yield indistinguishable patterns in summary statistics, leading to inherent non-identifiability (Frankel and Ané, 2026). Consequently, while find_graphs provides a flexible approach for estimating general network topologies, the inferred topology may not be uniquely determined when network complexity is high. Although find_graphs is commonly used to infer population histories, it has also been applied to interspecific and intergeneric relationships (Caldon et al., 2024; Owens et al., 2023). The *D*-statistic, a normalized version of *f*_4_ that forms the basis of find_graphs (Lipson, 2020), is also widely used to test for ancient gene flow at deep time scales, including among sea turtle (Vilaça et al., 2021), rattlesnakes (Myers et al., 2024) and angiosperms (Guo et al., 2021). Here, we assess the robustness of find_graphs to hidden paralogy and substitution rate variation at deep phylogenetic scale. SNaQ uses the frequency of gene trees as input, called quartet concordance factors (CFs) when pruned to a set of four taxa. The pseudolikelihood score combines the likelihood of these observed quartet CFs. PhyNEST uses site pattern frequencies computed from the sequence alignment for every 4-taxon set, and combines the likelihoods of these frequencies into a composite likelihood, assuming unlinked sites and a strict molecular clock (Kong et al., 2025). Unlike find_graphs, SNaQ and PhyNEST optimize their score over the restricted space of level-1 networks, which are identifiable from qCFs under the network coalescent model. However, this restriction limits their ability to resolve more complex reticulate histories (Pyron et al., 2025) (but see Kolbow et al., 2026, which lifts the level-1 restriction).

Like any other method, find_graphs, SNaQ, and PhyNEST rely on assumptions that may be violated in empirical datasets, motivating a systematic evaluation of their robustness to violations of model assumptions. In particular, many inference methods, whether based on summary statistics or explicit model fitting, implicitly assume relatively homogeneous substitution processes across genes and lineages (Cao et al., 2024), as well as the absence of gene duplication and loss (Kristensen et al., 2011; Shi, 2016). However, empirical datasets often deviate from these assumptions due to factors such as substitution rate heterogeneity across genes and lineages (Smith and Donoghue, 2008) and hidden paralogy. Hidden paralogy occurs when gene copies derived from duplication events are mistakenly treated as orthologs in phylogenetic analyses (Koonin, 2005; Siu-Ting et al., 2019). These sources of model misspecification can distort gene tree frequencies, site pattern frequencies, and summary patterns in the data, potentially leading to biased or incorrect network estimates (Lozano-Fernandez, 2022; Rasmussen and Kellis, 2012). Consequently, evaluating how sources of model violations influence phylogenetic network inference is essential for evaluating when we might trust an estimated species phylogeny (tree or network with reticulations), and when we might interpret them with caution. Here, we address this problem by focusing on two key sources of model violations: (1) substitution rate heterogeneity across genes and lineages, and (2) hidden paralogy arising from gene duplication and loss.

### 1.1. Substitution rate heterogeneity across genes and lineages

Variation in substitution rates is well-documented across genes and lineages (Duchêne et al., 2020; Smith and Donoghue, 2008; Thomas et al., 2010). Such heterogeneity can increase the likelihood of long-branch attraction (Bergsten, 2005; Bleidorn, 2017), thereby introducing discordance between estimated gene trees and the underlying species history. For example, in squamate reptiles, iguanians and snakes have been found to share faster rates of molecular evolution, which may cause long-branch attraction and misplacement of iguanians (Mongiardino Koch and Gauthier, 2018). Many commonly used methods for detecting introgression rely on assumptions of rate homogeneity or clock-like evolution, and violations of these assumptions can bias inference (Frankel and Ané, 2023, 2026; Koppetsch et al., 2024).

Focusing on methods based on biallelic patterns, Frankel and Ané (2023) showed that the D-statistic, D3 test, and HyDe are highly sensitive to rate variation, with lineage-specific rate heterogeneity leading to substantial inflation of false-positive rates of introgression. Pang et al. (2025) further showed that these site pattern tests fail even at shallow evolutionary depths, where small departures from clock-like rates are enough to produce false introgression signal in genome-scale data. This bias arises because rate variation generates asymmetric patterns of shared derived alleles through homoplasy, thereby mimicking signal of gene flow. Extending this result, Frankel and Ané (2026) demonstrated that the *f*_4_-statistic is likewise sensitive to violations of rate assumptions, fundamentally altering the interpretation of allele-frequency correlations. Koppetsch et al. (2024) further examined the impact of substitution rate variation on the *D*statistic, *D*_tree_ (Ronco et al., 2021), QuIBL (Edelman et al., 2019), SNaQ, and the MMS17 method (Meyer et al., 2017), a divergence-time-based introgression detection method. They found that homoplasy driven by rate heterogeneity can generate spurious introgression signal, with effects that are especially pronounced in deeper phylogenies and for the D-statistic, QuIBL and MMS17. *D*_tree_ and SNaQ showed no or little detectable sensitivity to rate variation. Their robustness can be explained by their use of gene tree topologies (without branch lengths) as input. Since rate variation primarily distorts branch lengths rather than tree topologies, topology-based inference is robust, although may be affected if rate variation is severe enough to induce systematic biases like long branch attraction. PhyNEST is a complementary approach because it uses site patterns under an equal substitution rate model (JC69) and a strict molecular clock. Building on previous evaluations of rate variation, we compare PhyNEST, SNaQ, and find_graphs on the same simulated data.

### 1.2. Hidden paralogy arising from gene duplication and loss

Another important violation of assumptions underlying phylogenetic network inference is paralogy arising from gene duplication and loss (Smith and Hahn, 2021). Gene duplication is pervasive (Campbell et al., 2019), particularly in plants, where up to *∼*65% of genes have duplicated copies (Panchy et al., 2016). In plants, whole-genome duplications occurred before the diversification of living angiosperms (Amborella Genome Project, 2013) and within water lilies and Chloranthales (Guo et al., 2021; Zhang et al., 2020b). Because paralogous sequences do not share a purely speciation-driven history, they can introduce systematic discordance between gene trees and the species tree (Martin and Burg, 2002; Xiong et al., 2022). A more insidious consequence is *hidden paralogy* (pseudo-orthology), where duplication followed by differential gene loss leaves a single copy per individual that is mistakenly treated as an ortholog (Smith and Hahn, 2021; Xiong et al., 2022). By distorting the distribution of gene tree topologies, hidden paralogy can bias species tree inference and, in some cases, lead to statistical inconsistency when misleading topologies are preferentially recovered across loci (Smith and Hahn, 2021; Xiong et al., 2022). Brown and Thomson (2017) demonstrated this in the context of reptile and turtle evolution: just two paralogous genes out of 248 in the transcriptome dataset of Chiari et al. (2012) were sufficient to flip the inferred placement of turtles entirely —from sister to archosaurs to sister to crocodilians— each with posterior probability of 1, in a Bayesian analysis of the concatenated alignment. This shows that even a negligibly small fraction of loci affected by hidden paralogy can produce strong, confident, and wholly incorrect phylogenetic conclusions, from some methods. This motivates us to conduct an evaluation of how hidden paralogy affects phylogenetic inference.

Two general strategies are used to handle paralogs: (1) filter loci to try and retain only single-copy orthologs, and (2) use methods that explicitly model duplication and loss. Most empirical phylogenomic pipelines follow the first approach (Joyce et al., 2025). Empirical studies in amphibians (Lissamphibia) show that removing paralogs can improve phylogenetic accuracy (Siu-Ting et al., 2019). This approach can filter out visible paralogous genes when several copies are recovered. Hidden paralogs, however, leave exactly one copy per species and survive any single-copy filter. In yeast for example, 4–7% of single-copy genes between any two species are hidden paralogs (Scannell et al., 2006).

Alternatively, several inference methods directly account for paralogy. For example, ASTRAL-multi (Rabiee et al., 2019) extends the ASTRAL algorithm to multi-copy genes and is statistically consistent under duplication-loss models (Legried et al., 2021; Markin and Eulenstein, 2021). ASTRAL-Pro (Zhang et al., 2020a) further improves performance by distinguishing duplication from speciation events to perform inference based on a restricted subset of speciation-consistent quartets. More broadly, quartet-based methods are robust to hidden paralogy because they rely on the dominant signal across gene tree quartets (Legried et al., 2021; Markin and Eulenstein, 2021; Smith and Hahn, 2021). However, this robustness is known for species tree inference only. Whether the presence of hidden paralogy would be misinterpreted as a signal of reticulation during network inference remains an open question.

The impact of hidden paralogy on phylogenetic inference depends on whether it alters gene tree topology, branch lengths, or both. Smith and Hahn (2022) modeled pseudoorthologs (hidden paralogy) on a rooted three-taxon species tree ((A,B),C) where a gene duplication occurs on the branch ancestral to all three taxa, and subsequent losses leave exactly one copy per species. They classified the outcomes into concordant and discordant pseudoorthologs. A concordant pseudoortholog arises when species A and B retain copies from the same paralog subclade, preserving the species tree topology but with a longer internal branch than a true ortholog. A discordant pseudoortholog arises when A and B retain paralogous copies through reciprocal loss; the resulting gene tree conflicts with the species tree, grouping either A or B with C depending on which copy C retains. Smith and Hahn (2022) proposed this classification primarily in a rooted 3-taxon setting. In an independent analysis, Xiong et al. (2022) showed that the timing of gene loss following whole-genome duplication determines the severity of the effect: when loss occurs soon after whole genome duplication, the retained copies are all from one paralog lineage and the gene tree matches the species tree. In contrast, loss along internal branches leads to differential paralog retention across lineages, producing pseudoorthologs that conflict with the species tree. Both kinds of pseudoorthologs are relevant to network inference. Discordant pseudoorthologs can alter the gene tree topology frequencies used by SNaQ, and the downstream pattern frequencies used by find_graphs and PhyNEST. Concordant pseudoorthologs alter branch lengths, affecting the pattern frequencies used by find_graphs and PhyNEST, and possibly the accuracy of estimated gene trees used by SNaQ.

Under the DLCoal model, the evolution of a gene family follows two steps (Rasmussen and Kellis, 2012). First, a birth-death process of duplication and loss generates a *locus tree*. Second, a *gene tree* is generated within each locus tree according to the coalescent process, thereby reflecting duplications, losses and ILS. Building on the discordance between the species tree and locus tree to focus on duplications and losses only, we distinguish three categories of hidden paralogy (Fig. 1).

1. **false paralogy:** duplications and losses occur but do not alter the topology or branch length of the locus tree for the specific retained copies (Fig. 1, top right). This corresponds to orthologs in the classification of Smith and Hahn (2022), where loss eliminates the original copy before a sampled speciation. We keep this category to quantify how often duplication and loss leave no trace in single-copy loci.
2. **Weak hidden paralogy:** the locus tree has the same topology as the species tree, but with distorted branch lengths (Fig. 1, bottom left). This corresponds to concordant pseudoorthologs, where the locus tree has longer internal branch(es) due to one or more split time in the locus tree corresponding to a duplication event, prior to the speciation time of the associated node in the species tree (Smith and Hahn, 2022).
3. **Strong hidden paralogy:** the locus tree topology conflicts with the species tree (Fig. 1, bottom right). This corresponds to discordant pseudoorthologs, arising from reciprocal paralog loss (Smith and Hahn, 2022), or from differential loss along internal branches following duplication of a locus or of the whole genome (Xiong et al., 2022).

**Figure 1.**
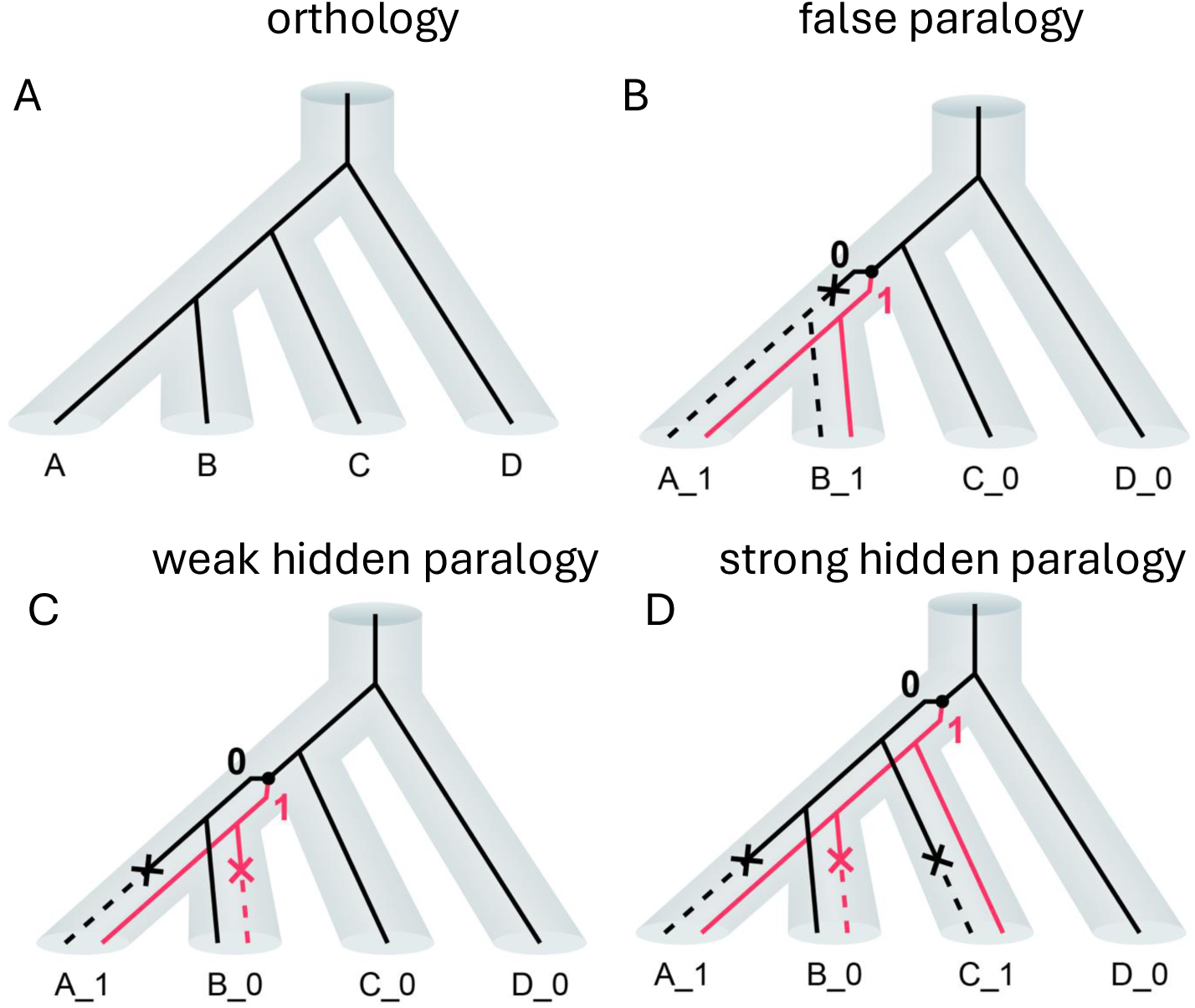
Conceptual illustration of orthology and paralogy. Top left: A possible species tree on taxa A-D (depicted as tubes) and a possible locus tree for one individual from each species (thin lines). Top right: Scenario illustrating false paralogy. A gene duplication event is followed by immediate loss of the original locus, such that the locus still has a topology identical to the species tree. This locus is then orthologous (splits occur at speciation events), but with duplications and losses happening during its evolutionary history. The duplication event can be summarized by a transposition to a new location in the genome, in A and B. The locus is indicated by a number in each sequence name, with 0 for the original locus and 1 for the duplicated location after the first duplication. Bottom left: Scenario illustrating weak hidden paralogy. Duplication and loss events occur such that there is at most one copy of the gene in each individual, and the locus tree topology matches the species tree, but branch lengths differ due to differential retention of paralogous copies. Bottom right: Scenario illustrating strong hidden paralogy. Duplication and differential losses result in retention of paralogous copies that generate a topologically discordant locus tree.

To evaluate whether substitution rate variation and/or hidden paralogy generate spurious signal of reticulation, we conducted a thorough simulation study using the three methods find_graphs, SNaQ and PhyNEST, with both violations calibrated from two empirical datasets. For each method, we measured how often it falsely favored reticulation, failed to recover the true species tree, and estimated a substantial proportion of gene flow.

## 2. Methods

### 2.1. Species tree for simulations

We simulated data on two eight-taxon species trees, referred to as the reptile and plant species trees. We calibrated the trees and simulation parameters using reptile UCE and angiosperm nuclear-gene datasets, respectively. For the reptile tree, we used a species tree and parameters estimated from 1,145 nuclear ultraconserved elements (UCEs) assembled by Crawford et al. (2012) to study the phylogenetic relationships of reptiles and turtles. According to Brown and Thomson (2017), who examined six datasets on reptile and turtle evolution, this UCE dataset had the lowest proportion (1.8%) of ‘spurious’ genes, defined as genes rejecting at least one well-established amniote relationship, compared with 6.8% to 22.1% in other datasets. They also found more homogeneous Bayes factor support across genes. This makes the UCE dataset a well-characterized empirical template for parameter estimation. The original dataset sampled two crocodilians, two birds, two turtles, one lizard, one snake, one tuatara, and human as the outgroup. For the plant species tree, we used the published alignments of the Angiosperms353 nuclear genes from the Kew Tree of Life Explorer, release 4.0 (Baker et al., 2022; Zuntini et al., 2024). We chose early-diverging angiosperms because their genomes document an ancient whole-genome duplication preceding angiosperm diversification (Amborella Genome Project, 2013), as well as subsequent duplications in water lilies and Chloranthales (Guo et al., 2021; Zhang et al., 2020b). Such duplications followed by differential gene loss can generate hidden paralogy, making these lineages an empirical template for understanding hidden paralogy. For the plant species tree, taxa A–H were *Pinus*, *Amborella*, *Nymphaea*, *Brasenia*, *Schisandra*, *Chloranthus*, *Liriodendron*, and *Ceratophyllum*, with the gymnosperm *Pinus* as outgroup. We extracted these samples from the published alignments, removed columns containing only gaps and ambiguous bases, and retained the 260 loci with sequence data across all selected taxa.

We used the same tree estimation workflow for both datasets. Gene trees were estimated separately for each locus with IQ-TREE (Minh et al., 2020), with substitution models selected by ModelFinder (Kalyaanamoorthy et al., 2017) and branch support assessed using 1,000 ultrafast bootstrap replicates (Hoang et al., 2018). The estimated gene trees were supplied to ASTRAL (Mirarab et al., 2014) for species tree estimation. For both trees, ASTRAL provided internal branch lengths *τ_i_* in coalescent units but did not estimate terminal branch lengths or the ingroup stem. To obtain input species trees for SimPhy, we assigned these missing lengths while retaining the estimated internal branch lengths. For the reptile tree, we assigned 0.5 coalescent units to the terminal branch of *Anolis* and to the ingroup stem. For the plant species tree, we assigned terminal branch lengths and the ingroup stem using empirical branch lengths *d_i_* in substitutions per site, as described below (Substitution Rate Variation). On all five internal branches, the ratios of coalescent to substitution lengths were similar (19.6–28.8 coalescent units per substitution per site), supporting a common conversion factor. We then chose the longest internal branch, leading to the common ancestor of *Nymphaea* and *Brasenia*, to calculate this factor (*k* = 3.77*/*0.172 *≈* 22 coalescent units per substitution per site). We averaged the substitution lengths of the *Nymphaea* and *Brasenia* terminal branches and multiplied this average by *k*, assigning 2.94 coalescent units to each branch. For the outgroup, sequence distances estimate the total length of the path from *Pinus* to the angiosperm crown, comprising the outgroup branch and ingroup stem. We converted this total to coalescent units using *k* and separated the two branch lengths by requiring all tips to be equidistant from the root, giving an ingroup stem of 1.56 coalescent units. For both trees, we then assigned the remaining terminal branch lengths using ultrametrize! from QuartetNetworkGoodnessFit.jl v.1.0.0.

As SimPhy requires branch lengths in generations, branch lengths for both trees were converted to numbers of generations by multiplying by a reference effective population size 2*N_e_*_,ref_ = 1000, yielding *t*_gen,*i*_ = 1000 *τ_i_* generations per branch. The choice of 2*N_e_*_,ref_ = 1000 is arbitrary: under the coalescent, gene tree distributions depend on the branch length in coalescent units, *τ_i_* = *t*_gen,*i*_ */*(2*N_e_*), so scaling branch durations and population sizes proportionally preserves gene tree distributions, up to a constant scaling of their branch lengths. The resulting plant tree was taller than the reptile tree (8.43 versus 3.44 coalescent units) and had a greater total length in generations. Because duplication and loss rates are specified per generation in SimPhy, lower duplication and loss rates were used for the plant tree, as described below. Both baseline trees were used as input to SimPhy to generate gene trees under duplication, loss, and incomplete lineage sorting.

### 2.2. Gene tree simulation

We simulated gene trees using SimPhy version 1.0.2 (Mallo et al., 2016) with each species tree as input. We varied four factors and used all their combinations, for a total of 3 *×* 3 *×* 2 *×* 2 = 36 parameter settings per species tree (Table 1). We applied find_graphs and SNaQ to all 36 settings. PhyNEST was applied to the 18 settings with one individual per taxon only, because it does not implement a mapping of multiple sampled individuals to their species. For each parameter combination, we simulated 100 independent replicates. All random seeds were derived deterministically from a hierarchical seed system based on the parameter configuration name, ensuring fully reproducible results across all stochastic components. The four simulation factors are described below in turn.

**Table 1.** Simulation parameters. Combining the four factors yields 3 *×* 3 *×* 2 *×* 2 = 36 settings. For each, 100 replicates were simulated, with 1000 genes per replicate. PhyNEST was run on the 18 settings with *n* = 1 individual per taxon.

| **Factor** | **Levels** |
|---|---|
| Substitution rate variation | none (N), gene-specific (G), lineage-specific (L) |
| Duplication and loss rate | 0, 3 × 10<sup>−4</sup>, 4 × 10<sup>−4</sup> (reptile)<br>0, 1 × 10<sup>−4</sup>, 1.4 × 10<sup>−4</sup> (plant) |
| ILS level | *N<sub>e</sub>* = 500 (low ILS), *N<sub>e</sub>* = 1000 (high ILS) |
| Individuals per taxon | *n* = 1, *n* = 2 |

#### Incomplete Lineage Sorting

ILS was controlled via SimPhy’s -sp parameter, which sets the haploid effective population size (*N_e_*) applied uniformly across all branches. Setting -sp 1000 matches the reference population size (2*N_e_*_,ref_ = 1000), so the effective branch lengths in coalescent units equal the empirical *τ_i_* shown in Fig. 2. This constitutes our high-ILS setting. Setting -sp 500 halves the population size relative to the reference, doubling the effective branch lengths in coalescent units and decreasing the probability of deep coalescence. This constitutes our low-ILS setting.

**Figure 2.**
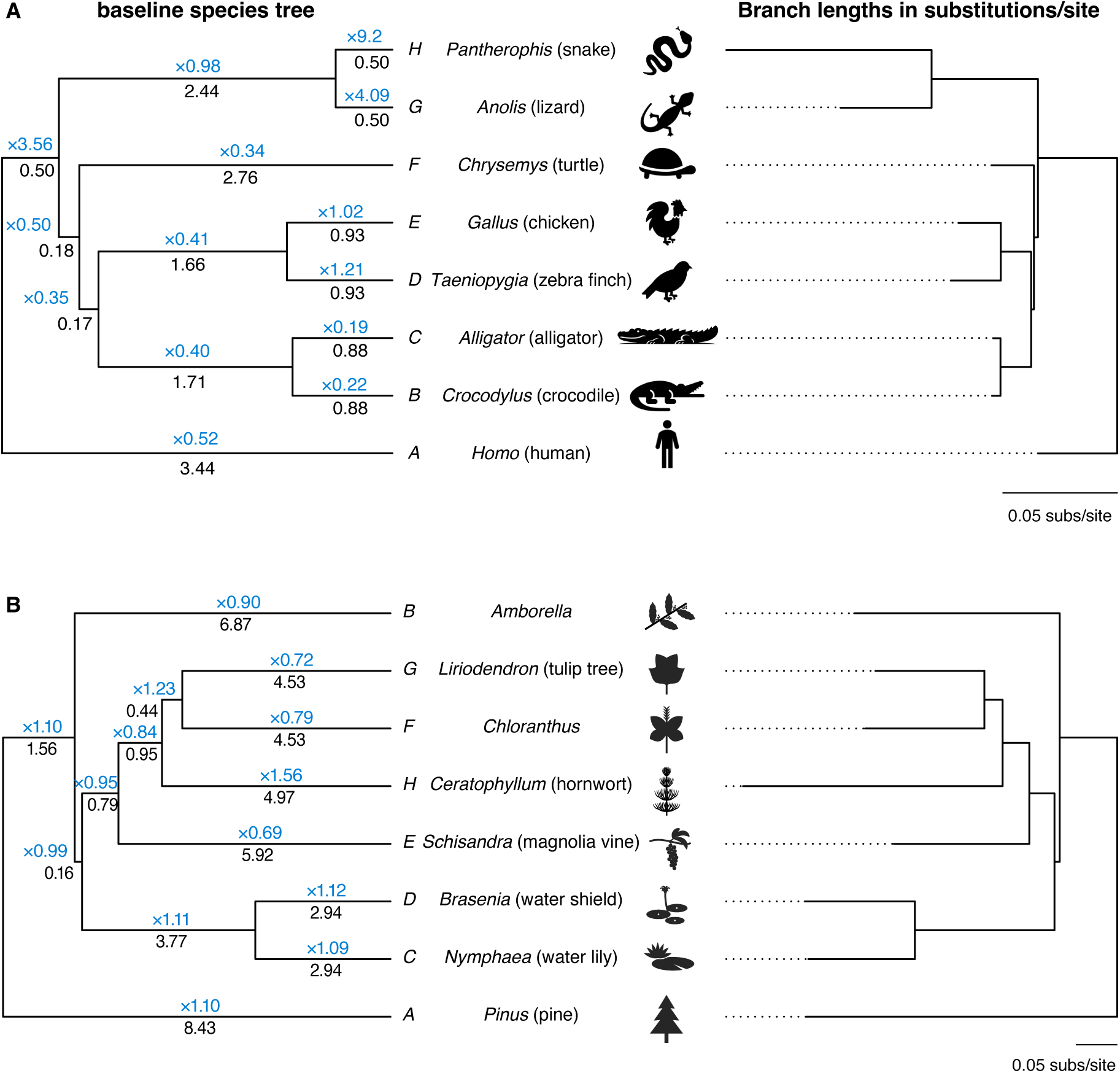
Baseline species trees and lineage-multiplied species trees used in simulations. (A) The baseline reptile species tree for eight taxa: *Homo* (human, A), *Crocodylus* (crocodile, B), *Alligator* (alligator, C), *Taeniopygia* (zebra finch, D), *Gallus* (chicken, E), *Chrysemys* (turtle, F), *Anolis* (lizard, G), and *Pantherophis* (snake, H). (B) The baseline plant species tree for *Pinus* (A, outgroup), *Amborella* (B), *Nymphaea* (C), *Brasenia* (D), *Schisandra* (E), *Chloranthus* (F), *Liriodendron* (G), and *Ceratophyllum* (H). Black numbers on the left: length *τ_i_* in coalescent units for branch *i*. Under a constant substitution rate, the expected substitution length per branch is *τ_i_ r*, where *r* = 0.0195 for reptiles and 0.0412 for plants, in substitutions per site per coalescent unit. Blue numbers are lineage-specific rate multipliers, *m_i_* for branch *i* (see methods for details). Right: The corresponding trees, with branch lengths scaled proportional to *τ_i_ m_i_*, and scaled by the average genome-wide rate *r* to show the expected substitution accumulation per branch under lineage-specific rates. Scale bars: 0.05 substitutions per site.

#### Number of individuals

Empirical phylogenomic datasets routinely include multiple individuals per species or population to capture within-population genetic diversity and improve allele-frequency estimation. For this reason, we simulated two sampling depths: one individual per taxon and two individuals per taxon. We ran find_graphs and SNaQ at both depths, and PhyNEST at one individual per taxon only, as explained above. This number was controlled via the SimPhy parameter -si. When two individuals were simulated per taxon, the downstream pipeline was adjusted to estimate phylogenies at the taxon level.

#### Substitution Rate Variation

To generate realistic rate variation, we analyzed the variation in gene trees across loci from the reptile and the plant dataset, with branch lengths in substitutions per site. We applied the following procedure separately to each dataset. First, for each locus *l*, an outgroup-to-ingroup sequence divergence for this locus was estimated by the height of its IQ-TREE-inferred gene tree, 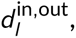 taken as the median genetic distance between the in-group taxa and the outgroup taxon (*Homo* for reptiles and *Pinus* for plants). The median was chosen for robustness and to limit the influence of lineages with extreme rates. A small vs. large 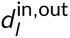 value reflects a slow-vs. fast-evolving locus. The average 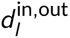 across the 1,145 reptile and 260 plant loci was 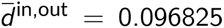 and 0.7173, respectively. The smaller sequence divergence in the reptile dataset reflects the UCE’s marker conservation by design. Second, for each locus *l*, its gene tree was rescaled by 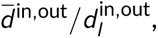 where 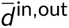 is the average for that dataset given previously, to account for rate variation across loci, before calculating the average genetic distance between each pair of taxa, as the average across gene trees. Finally, these pairwise distances were fitted to the species tree topology using ordinary least-squares with calibratefrompairwisedistances! from PhyloNetworks, to obtain branch lengths *d_i_*, in substitutions per site, that best represent the observed genetic distances between taxa (Fig. 2 right). *No variation*: To generate data under a molecular clock without rate variation, we used an average genome-wide rate 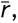 obtained separately for each species tree as the ratio of total tree length in substitutions per site to coalescent units:

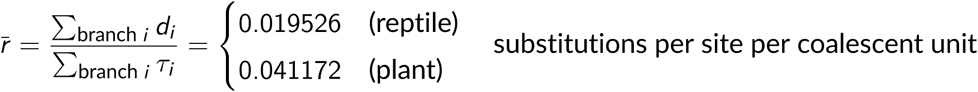

where *τ_i_* is the length of branch *i* in coalescent units as described previously (shown in Fig. 2 left). Given that the number of generations for each branch in the species tree was set based on the haploid effective population size *N_e_*_,ref_, the per-generation substitution rate was set to

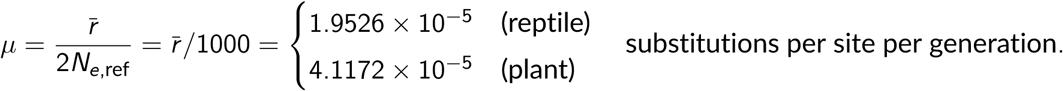

This rate *µ* was fixed across all simulation settings via SimPhy’s -su parameter. Without any additional multipliers, this setting corresponds to no rate variation.

#### Rate variation across genes

For each gene family, a rate multiplier *r_g_* was drawn independently from a log-normal distribution with log-scale mean *a* and standard deviation *σ*: ln(*r_g_*) *∼ N* (*a*, *σ*^2^) with (*a*, *σ*) *≈* (*−*0.19, 0.6164) or (*−*0.03634, 0.2696) for the reptile and plant tree respectively. These distributions were implemented via SimPhy’s option -hl, using the corresponding parameters for each species tree. These standard deviation were estimated by fitting a log-normal distribution to the normalized tree heights 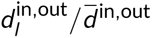 for each dataset, obtained as described earlier. The distribution of these gene tree heights was fitted to a log-normal distribution, and also to a normal and Gamma distribution. The log-normal distribution provided a better fit for both datasets. The mean parameter was set to *a* = *−σ*^2^*/*2 and was chosen to obtain relative rate multipliers with mean 1 across loci, as

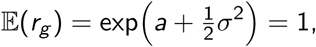

so the mean substitution rate across loci, E(*r_g_µ*), remains equal to the genome-wide *µ*.

#### Rate variation across lineages

For each species tree, each branch *i* was assigned a lineage-specific multiplier

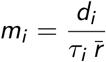

where *d_i_* is the empirical substitution length of branch *i* described above (see Fig. 2 right). Specifically, each branch *i* in the SimPhy species tree was specified as generations_i_\**m_i_*, where SimPhy interprets the multiplier *m_i_* as scaling the baseline substitution rate *µ* (from the -su option) for that branch alone. Combined with baseline rate *µ*, the expected number of substitutions per site along branch *i* was then

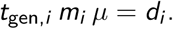

By using the empirical substitution length *d_i_* for each branch *i* (Fig. 2 right), we recover the empirical pattern of rate variation across lineages for each empirical dataset. For example, in the reptile data, *Pantherophis* (snake, H) accumulates substitutions far faster than other lineages (Fig. 2 A). In the plant data, *Ceratophyllum* had the highest terminal-branch substitution rate. (Fig. 2 B).

Duplication and loss rates. To simulate hidden paralogy, we used the DLCoal (Duplication, Loss, and Coalescence) model from Rasmussen and Kellis (2012), implemented in SimPhy. We imposed equal gene duplication and loss rates using SimPhy’s -lb and -ld flags, set to 0 (no paralogy), 3 *×* 10*^−^*^4^ or 4 *×* 10*^−^*^4^ per locus per generation on the reptile species tree, and 0, 1 *×* 10*^−^*^4^ or 1.4 *×* 10*^−^*^4^ on the plant tree. Equal duplication and loss rates maximize the probability that a gene tree retains exactly one surviving copy per taxon after gene loss, making paralogous copies appear single-copy despite their duplication history. We required a minimum of eight tips per locus tree (-ll 8), matching the total number of sampled taxa. The three duplication and loss rate settings correspond to 0%, *∼*4–5%, and *∼*12% of retained simulated gene trees exhibiting strong hidden paralogy, respectively, for both species trees (Table 2).

**Table 2.** Summary of the percentage of gene trees that are orthologs (without any duplication/loss events) and that are paralogs of different hidden paralogy (HP) types, for the reptile and the plant trees. Percentages are aggregated across levels of ILS, substitution rate variation, and number of individuals per taxon, and are reported as mean *±* standard deviation. The duplication/loss rate is per locus per generation. With duplication and loss rates set to zero, both trees yielded 100% orthologs and 0% in all HP categories, with no variation across settings.

| duplication/<br>loss rate | species<br>tree | rate | orthologs<br>(%) | false paralogy<br>(%) | weak HP<br>(%) | strong HP<br>(%) |
|---|---|---|---|---|---|---|
| medium | reptile | 3 × 10<sup>−4</sup> | 8.7 ± 0.08 | 76.0 ± 0.14 | 9.8 ± 0.13 | 5.5 ± 0.07 |
|  | plant | 1 × 10<sup>−4</sup> | 13.5 ± 0.08 | 76.9 ± 0.05 | 5.4 ± 0.06 | 4.2 ± 0.07 |
| high | reptile | 4 × 10<sup>−4</sup> | 1.7 ± 0.04 | 69.7 ± 0.20 | 16.6 ± 0.10 | 12.0 ± 0.12 |
|  | plant | 1.4 × 10<sup>−4</sup> | 2.6 ± 0.05 | 74.6 ± 0.13 | 10.6 ± 0.08 | 12.3 ± 0.12 |

Gene tree filtering. To systematically simulate genes with hidden paralogy, we applied an approach to iteratively simulate gene trees while filtering invalid gene trees to reach the target number of *N* = 1000 gene trees. After each SimPhy batch, we discarded gene trees in which any taxon appeared more than once (detectable paralogy) or any of the eight taxa was missing after gene loss. Allowing missing taxa would cause missing genotypes in the downstream simulated sequence alignments and the EIGENSTRAT files, introducing bias when computing *f*_2_ statistics later used for find_graphs (Maier et al., 2023). If fewer than 1,000 were retained, SimPhy was re-run iteratively up to 20,000 iterations (Fig A1 in the Supplementary Material, SM). Under the two lower duplication–loss rates, the target of 1,000 genes was almost always achieved, except for four reptile replicates under medium duplication rate, which reached the iteration limit with 999 genes. Under the high duplication rate, every replicate reached the iteration limit: the mean number of retained gene trees was 958 (range 940–977) for the reptile species tree and 947 (range 928–966) for the plant species tree.

### 2.3. Simulation of Molecular Sequences

Filtered gene trees were passed to SeqGen v.1.3.4 (Rambaut and Grassly, 1997) to simulate DNA alignments of 1,000 base pairs per locus under the HKY+Γ model (Hasegawa et al., 1985). To capture realistic among-locus heterogeneity, we first extracted the parameters of the substitution models estimated for the empirical loci in each dataset, then used their distribution across loci, fitted by a parametric family, to simulate parameters for each gene (Table 3). The HKY rate ratio *κ* was estimated from loci whose selected model was HKY. For each simulated locus, the sampled *κ* was converted using its base frequencies to the transition/transversion ratio required by Seq-Gen’s -t option. The distribution of the equilibrium base frequencies and of the Gamma distribution’s shape *α* to simulate rate variation across sites were taken from loci whose selected models included these parameters. For *κ* and *α*, we fitted normal, log-normal, and Gamma distributions and selected the family with the highest log likelihood. Base frequencies were fitted to a Dirichlet distribution.

**Table 3.**
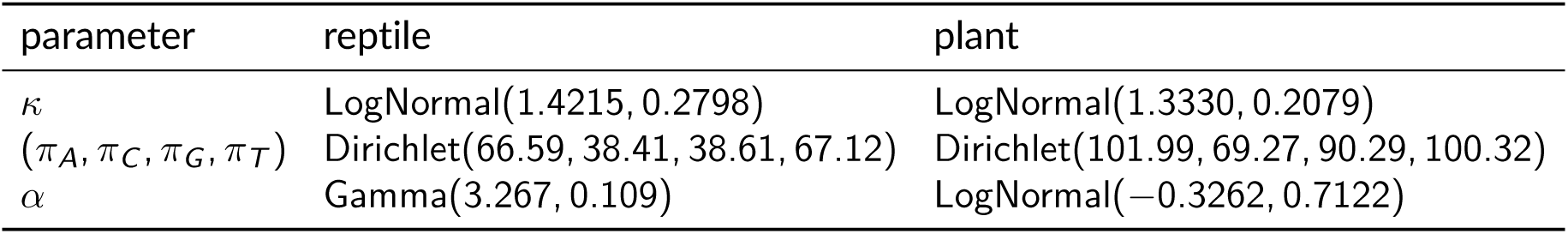
Distribution of the substitution model parameters across genes. The HKY model uses a transition/transversion rate ratio *κ* and base frequencies. Rate variation across sites (within a gene) was simulated with a Gamma distribution of mean 1 and shape *α*. Log-normal parameters are log-scale means and standard deviations.

| parameter | reptile | plant |
|---|---|---|
| κ | LogNormal(1.4215, 0.2798) | LogNormal(1.3330, 0.2079) |
| (π<sub>A</sub>, π<sub>C</sub>, π<sub>G</sub>, π<sub>T</sub>) | Dirichlet(66.59, 38.41, 38.61, 67.12) | Dirichlet(101.99, 69.27, 90.29, 100.32) |
| α | Gamma(3.267, 0.109) | LogNormal(−0.3262, 0.7122) |

For later analysis with find_graphs, the sequence alignments of all genes from each replicate were concatenated into a single FASTA file (one per replicate). This concatenated alignment was converted to VCF using snp-sites v.2.5.1 (Page et al., 2016), then to EIGENSTRAT format using the vcf2eigenstrat.py script (https://github.com/mathii/gdc), keeping biallelic sites only.

Each haploid sequence was coded by whether it carried the allele of the outgroup species of taxon A.

### 2.4. Estimation of gene trees and species tree

Gene tree estimation. For each simulated sequence alignment, a gene tree was estimated using IQ-TREE v2.4.0 (Minh et al., 2020) under the HKY+Γ model (-m HKY+G), with base frequencies estimated empirically from each alignment (the IQ-TREE default). Estimated gene trees within each replicate were used for calculating concordance factors, which served as input to SNaQ (described below).

Gene tree variability. To decompose the variability of estimated gene trees into distinct contributions from ILS, hidden paralogy and estimation error, we computed two normalized Robinson-Foulds (RF) distances for each locus. First, the gene tree discordance was measured as the RF distance between the simulated (true) gene tree output from SimPhy and the species tree topology. This topological conflict quantifies the contribution of ILS for orthologous and for false and weak HP loci, and both contributions of ILS and hidden paralogy for strong HP loci. Second, the gene tree estimation error was measured as the RF distance between the simulated (true) gene trees and the gene trees estimated by IQ-TREE to quantify estimation error introduced by sequence-based tree inference. Both distances were normalized by 2(*m −* 3), where *m* is the number of tips in the trees being compared, yielding values between 0 and 1. For gene tree estimation error, *m* = 8*n*_ind_ sampled individuals with *n*_ind_ = 1 or 2 per species. For gene tree – species tree discordance, we used *m* = 8 taxa. To compare the species tree to a gene tree *G* with two individuals per species, we averaged the RF distance to each subtree of *G* obtained by retaining one individual per species, then averaged these distances. These distances are summarized in Appendix B (Table B1). All RF distances were computed using mudistance_semidirected in PhyloNetworks.jl v.1.1.0 (Solís-Lemus et al., 2017).

Species tree estimation. For each replicate, a species tree was obtained from the estimated gene trees using ASTRAL-IV v1.24.4.8 (Zhang et al., 2025). For replicates with two individuals per taxon, a mapping file specifying which accessions belonged to which species was supplied to ASTRAL. Although ASTRAL-Pro (Zhang et al., 2020a) is intentionally designed to handle paralogs, it can only detect gene duplication with multiple gene copies and yield the same topology as ASTRAL in single-gene-copy datasets. We used ASTRAL, as it is recommended for single-gene-copy dataset by the developers (Zhang et al., 2025) for its ability of handling polytomies and different algorithms of calculating branch support. The resulting species tree served as the starting topology for network estimation with SNaQ and PhyNEST.

### 2.5. Network estimation

Phylogenetic networks were estimated under a tree model without reticulation (*h* = 0) and a network model with at most one reticulation (*h* = 1) using find_graphs (Maier et al., 2023), SNaQ (Solís-Lemus and Ané, 2016) and PhyNEST (Kong et al., 2025). Model selection between *h* = 0 and *h* = 1 was based on *f* -statistic residuals for find_graphs, a quartet goodness-of-fit test for SNaQ, and a calibrated difference in per-site composite-likelihood scores for PhyNEST. find_graphs.. For each replicate, the concatenated alignment in EIGENSTRAT format was used as input to compute block-jackknifed *f*_2_ statistics with f2_from_geno using adjust_pseudohaploid=TRUE, since each simulated individual carried a single haploid sequence rather than a diploid genotype. The block size was set to 1,000 sites (blgsize=1000) to match the simulated gene length, so that each jackknife block corresponds to one gene. For each replicate, 100 independent find_graphs searches were performed under both the tree model (*h* = 0) and the single-reticulation model (*h* = 1), using taxon A as the outgroup and the software-default of 100 search generations per run (stop_gen=100). The multiple-run strategy was adopted to mitigate local optima, consistent with the developer recommendation (Maier et al., 2023).

Within each replicate and separately for *h* = 0 and *h* = 1, only graphs within two score units of the best-scoring graph across all runs were retained. Some graph topologies were duplicated, if they were found in multiple runs. Such duplicates were removed, keeping unique topologies. Since the find_graphs score is equal to the negative likelihood, up to some constant, a lower score indicates a better fit. This threshold corresponds to ΔAIC = 4 for models of equal complexity. Burnham and Anderson (2004) interpret models with ΔAIC *≤* 2 as having “substantial” support, those with 4 *≤* ΔAIC *≤* 7 have“considerably less” support. Our threshold sits at the boundary between these two categories. This excludes weakly supported models while retaining multiple graphs as recommended by Maier et al. (2023).

Each retained graph, as well as the true graph, was then evaluated using qpgraph to compute its worst absolute *f* -statistic residual *z* -score, traditionally called worst residual (WR). This quantity summarizes the largest discrepancy between observed and graph-predicted *f* -statistics across all 406 distinct contrasts for a topology with eight taxa (residuals from 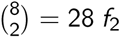 values, 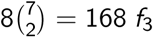 values, and 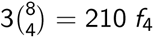 values). We evaluated the goodness-of-fit of each graph using two WR thresholds: the standard WR *≤* 3.0 criterion (Maier et al., 2023) and a more permissive threshold of WR *≤* 3.7 to classify the graph as providing a good fit to the data. The latter was motivated by the empirical 95^th^ percentile of the WR values on the true graph topology in our simulations (see Results). Under each threshold, the tree model (*h* = 0) was accepted if any retained tree (graph with *h* = 0) satisfied the criterion; a single reticulation (*h* = 1) was inferred if any *h* = 1 graph satisfied the criterion while no *h* = 0 graph did; and *h >* 1 was concluded if no graph under either model satisfied the criterion.

SNaQ.. Quartet concordance factors (qCFs) were estimated from the estimated gene trees for each replicate using countquartetsintrees from PhyloNetworks v.1.1.0 (Solís-Lemus et al., 2017). When one individual was sampled per taxon, qCFs were computed directly at the individual level; when two individuals were sampled per taxon, qCFs were first computed at the individual level, and then species-level qCFs were obtained by using mapallelesCFtable in SNaQ.jl.

For each replicate, we performed 10 independent SNaQ optimization runs under the tree model (*h* = 0, hmax=0), initialized from the ASTRAL-estimated species tree. Ten runs is typically sufficient when searching for a tree (*h* = 0). We then performed 100 independent runs under the single-reticulation network model (*h* = 1, hmax=1), initialized from the *h* = 0 best result. Starting *h* = 1 runs from the *h* = 0 optimum is consistent with best practice recommended by the SNaQ developers and avoids poor initializations in network space (Solís-Lemus and Ané, 2016).

To assess model fit, we applied the goodness-of-fit test by Cai and Ané (2021) implemented in QuartetNetworkGoodnessFit.jl v.1.0.0, which compares observed quartet concordance factors with those expected under the multispecies coalescent on the candidate species phylogeny, using 1,000 parametric simulations to compute a *p*-value on the inferred tree (*h* = 0) and single-reticulation network (*h* = 1). The tree model (*h* = 0) was selected as a good fit to the data when the tree was not rejected (*p_h_*_=0_ *>* 0.05). Otherwise, a single-reticulation network (*h* = 1) was selected when *h* = 1 was not rejected but *h* = 0 was (*p_h_*_=1_ *>* 0.05 and *p_h_*_=0_ *≤* 0.05). Finally, when neither *h* = 0 nor *h* = 1 provided an adequate fit (*p_h_*_=0_ *≤* 0.05 and *p_h_*_=1_ *≤* 0.05), this outcome was recorded as *h >* 1 being favored.

PhyNEST.. Each concatenated alignment was analyzed with PhyNEST.jl v0.1.12 (Kong et al., 2025). For each set of four taxa, PhyNEST counts how often each site pattern occurs across the alignment and computes the probability of each pattern under the network multispecies coalescent. The likelihood of these counts is then multiplied over all four-taxon sets, as if they were independent, to form a composite likelihood, which is maximized over networks. These site-pattern probabilities assume the JC69 substitution model with equal rates across sites and a strict molecular clock (Kong et al., 2025). Our data therefore violated the substitution model of PhyNEST in all settings, as typical of empirical data, and the clock assumption under lineage-specific rates.

For each replicate, the concatenated alignment was converted to sequential PHYLIP format and used as input. Because PhyNEST assumes a single sampled lineage per species, settings with two individuals per species were dropped. For each replicate, we performed 100 independent search runs under both the tree model (*h* = 0, hmax=0), initialized from the ASTRAL-estimated species tree rooted at taxon A and under the single-reticulation model (*h* = 1, hmax=1), initialized from the best *h* = 0 topology, as for SNaQ. All other search settings were left at their software defaults.

PhyNEST returns the best-scoring network found with at most *h* reticulations as specified by the user, without a criterion to decide whether an additional reticulation is warranted (Kong et al., 2025). We therefore designed a model selection procedure for this study. The score reported by PhyNEST is the negative log composite likelihood of the estimated network, so a lower score indicates a better fit. We compared the two models with the per-site score difference

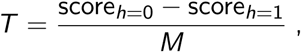

where *M* is the number of sites in the concatenated alignment of the replicate. We divided by *M* because the composite likelihood sums over all sites and over all 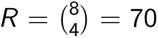 four-taxon sets, so the raw score grows linearly with *M* and with *R*. *R* was the same in all replicates, whereas *M* varied. In our study *M* is 1,000 sites times the number of retained loci, varied when filtering retained fewer than 1,000 loci (between 928 and 1000, see Gene tree filtering). Each replicate’s *T* was thus normalized by its own concatenated alignment length *M*.

Because the composite likelihood combines contributions from overlapping quartets, the usual *χ*^2^ reference distribution for a likelihood-ratio test does not generally apply (Varin et al., 2011). We therefore calculated a significance threshold *T^∗^* separately for each species tree and ILS level. Within each group, we pooled the *T* values across simulations with constant or gene-specific rates and all duplication and loss rates, then defined *T^∗^* as the 95th percentile of this distribution (SM Table C2). We selected the model with one reticulation when *T* exceeded this threshold *T^∗^* for the corresponding species tree and ILS level.

### 2.6. Evaluation metrics

Performance of network estimation was evaluated using three metrics.

Type I error rate.. Because the true species phylogeny is a tree, rejecting the tree model constitutes a type I error. The type I error rate was computed as the proportion of replicates in which *h* = 0 was rejected. We also recorded the proportions of replicates favoring *h* = 0, *h* = 1 or *h >* 1. For PhyNEST, model selection lead to favoring either *h* = 0 or *h ≥* 1. Only its type I error rate is reported (proportion of replicates rejecting *h* = 0, with *T > T^∗^*).

True tree recovery.. We evaluated whether the true species tree topology was recovered. Under *h* = 0, recovery corresponds to inferring the correct tree topology exactly. Under *h* = 1, the inferred network was classified as recovering the true tree if that tree was displayed in the network, that is, if the true tree could be obtained by deleting one hybrid edge from the network. All trees (true and displayed in estimated networks) were considered as unrooted. For SNaQ and PhyNEST, which return a single best-scoring topology per replicate, the inferred tree (*h* = 0) and network (*h* = 1) were each directly examined for true topology recovery. For find_graphs, from which we kept multiple candidate topologies per replicate, we evaluated topological accuracy in two ways: (1) whether the true tree was displayed in at least one of the retained graphs, and (2) whether the true tree was displayed in the single best-scoring graph.

In the reptile species tree, taxon F (turtle), recognized as difficult to resolve within amniotes (Chiari et al., 2012), occupies an interior position connected by two particularly short branches (0.17 and 0.18 coalescence units, Fig 2), causing true discordance about its position in gene trees, and making its placement difficult to resolve correctly. In the plant species tree, taxon B (*Amborella*), recognized as difficult to resolve at the base of angiosperms (Xi et al., 2014), is separated from the (C,D) clade by the shortest branch (0.16 coalescence units, Fig 2). We therefore additionally evaluated recovery after pruning the corresponding taxon (F for reptiles and B for plants) from both true and inferred topologies.

Estimated gene flow proportion.. For networks inferred under the single-reticulation model (*h* = 1), we recorded the minor inheritance probability *γ* (below 0.5), which quantifies the estimated proportion of gene flow contributed by the minor parental lineage. Under the null hypothesis of no true reticulation, elevated *γ* values near 0.5 indicate spurious signal of strong gene flow, whereas small values reflect weak or noise-driven reticulation signal.

## 3. Results

Our simulations generated datasets spanning a gradient from no paralogy to relatively high levels across the hidden paralogy (HP) categories (Table 2). As duplication and loss rates increased, the proportion of orthologous loci without any duplication or loss events declined to 8.7% and 1.7% in the reptile simulations and to 13.5% and 2.6% in the plant simulations, at the medium and high rates, respectively. The vast majority of loci were false paralogs, with a duplication and loss history but whose locus tree matched the species tree exactly (76.0% and 69.7% for the reptile simulations, and 76.9% and 74.6% for the plant simulations, at the medium and high rates, respectively). At the high duplication rate, the proportion of loci with strong HP was similar in the reptile and plant simulations (12.0% and 12.3%, respectively). Such a level of contamination would be considered substantial in empirical phylogenomic studies. Despite this variation in hidden paralogy levels, ASTRAL consistently recovered the true species tree in all replicates across all simulation settings.

### 3.1. Gene tree discordance and estimation error

The true gene trees of orthologous and weak HP loci differed from the species tree, due to ILS, by more than 1 out of 5 internal branches on average under low ILS (mean normalized RF distances of 0.21–0.24) and nearly 2 out of 5 branches under high ILS (0.33–0.38; see SM Fig. B1 and SM Table B1). Due to additional discordance from duplications and losses, gene trees with strong HP loci were more discordant than orthologous or weak HP loci. Across medium and high duplication rates, the mean distances of loci with strong HP were 0.36 and 0.40 for reptiles and 0.49 and 0.55 for plants (SM Fig. B1), showing that discordance at strong HP loci was substantially greater in the plant simulations. In our 8-taxon species trees, a 0.40 normalized RF distance corresponds to an average of 2 discordant branches out of the 5 internal branches. For example, among strong HP loci, at least three of the five internal branches differed from the reptile species tree in 16% *−* 25% of gene trees, compared with 46% *−* 60% of plant gene trees (SM Table B1).

Gene tree estimation error, as measured by the average normalized RF distance between estimated and true gene trees, was lower with two individuals per taxon than one. At both sampling depths, gene tree estimation error was highest under gene-specific rates, reaching about 0.20 with one individual per taxon (SM Fig. B1), corresponding to an average of 1 wrong branch out of 5. Overall, gene tree estimation error was moderate. Across species trees and rate variation settings, the estimated gene tree matched the true topology exactly in 37%–69% of cases with one individual per taxon and 30%–44% with two. Estimated trees that shared none of their internal branches with the true tree were rare and occurred exclusively under gene-specific rates (SM Table B2).

### 3.2. Empirical model-selection calibration

After network estimation with find_graphs, the standard criterion for assessing graph fit is to check that the worst absolute *f* -residual *z* -score (worst residuals, WR) falls below a threshold (Maier et al., 2023). To evaluate whether the commonly used threshold (WR *≤* 3) is appropriate under our simulation conditions, we computed the WR of the true species tree by fitting its branch lengths to the *f* statistics. For each species tree, we pooled WR values across duplication and loss rates, ILS levels, and numbers of individuals per taxon, for constant and gene-specific rates. The 95th percentiles exceeded 3.0 under constant rates (3.56 for the reptile tree and 3.41 for the plant species tree) and under gene-specific rates (3.75 and 3.49, respectively). Pooling the constant and gene-specific settings gave 95th percentiles of 3.65 for the reptile tree and 3.47 for the plant species tree. We therefore adopted the more permissive threshold WR *≤* 3.7 for both species trees: This was the smallest threshold, to one decimal place, exceeded by fewer than 5% of the true-tree WR values pooled across constant and gene-specific rates for each species tree. Under lineage-specific rates, WRs were higher under low ILS, so we pooled WR values separately for each ILS level. The 95th percentiles exceeded the cutoff of 3.7 for both species trees at both ILS levels and were much higher for the plant species tree (5.5 and 6.8 for the reptile tree and 16.8 and 21.7 for the plant species tree under high and low ILS, respectively; Fig. 3A, SM Table C1).

**Figure 3.**
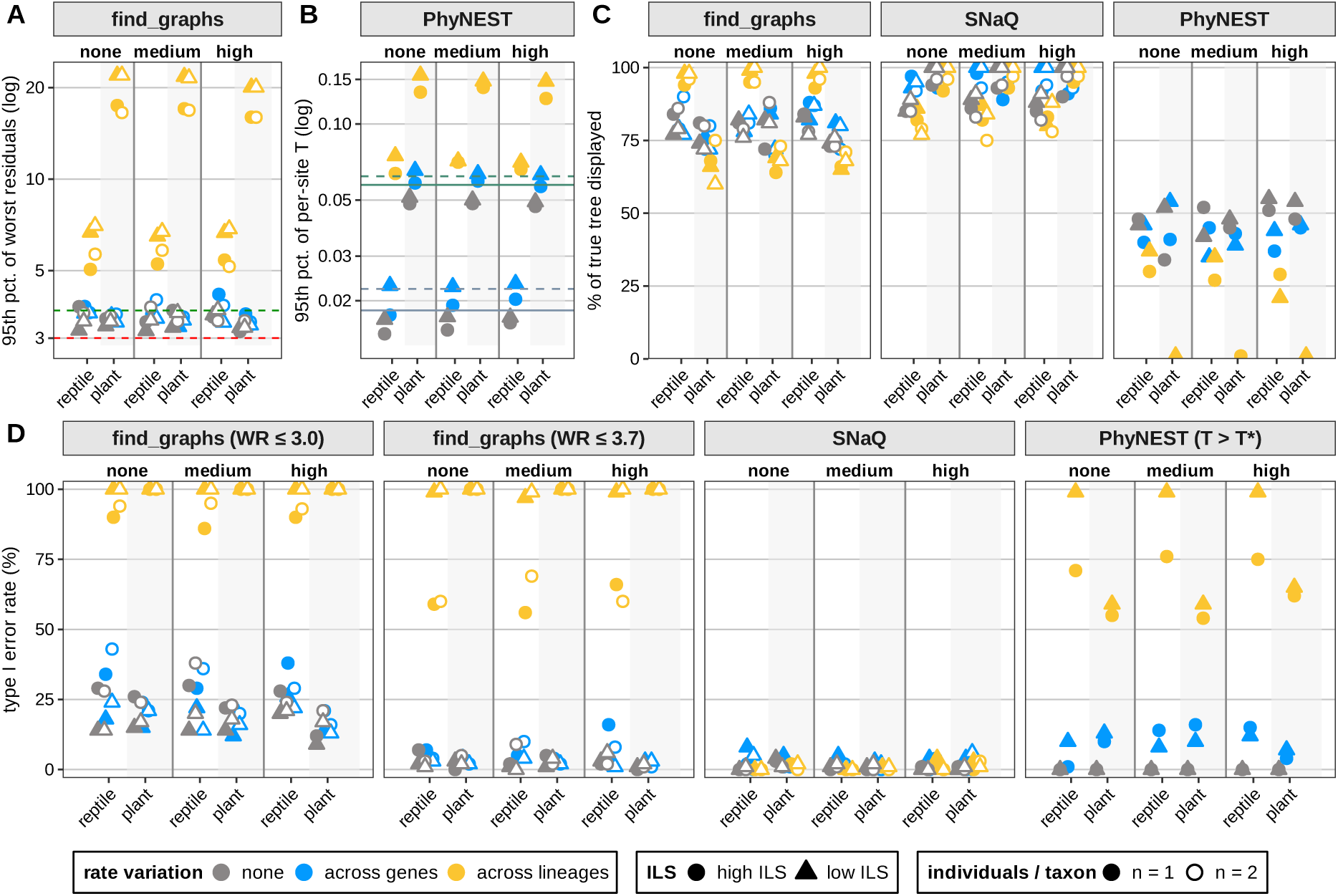
(A) 95th percentile of the worst residual (WR) for the true species tree fitted with find_graphs. Dashed lines mark WR thresholds of 3.0 (red) and 3.7 (green). (B) 95th percentile of the per-site score difference *T* from PhyNEST. Horizontal lines mark *T^∗^* for reptiles (grey) and plants (green), with solid and dashed lines for high and low ILS, respectively. Panels A and B use log scales. (C) Probability that the inferred network (single best-scoring graph for find_graphs) displays the true species tree under *h* = 1. (D) Type I error rates under the indicated model-selection criterion. Each point represents one simulation setting, with reptile and plant results paired within each duplication/loss level. Colors indicate substitution rate variation, shapes indicate ILS (circles, high; triangles, low), and filled and open symbols indicate one and two individuals per taxon, respectively. PhyNEST was run with one individual per taxon only.

For PhyNEST, we likewise used simulated datasets with constant and gene-specific rates to calibrate a model-selection threshold on the per-site score difference *T* . The 95th percentiles of *T* were overall higher under gene-specific rates for both species trees compared to constant rates, but these percentiles were much closer to one another than to the percentile under lineage-specific rates (SM Table C2). We therefore combined the constant and gene-specific rate settings for calibration. Under these rates, *T* varied little with duplication and loss levels but differed between ILS levels, so we pooled *T* separately for each species tree and ILS level. The 95th percentile of each distribution defined the threshold *T^∗^*. At low and high ILS, *T^∗^* was 0.022 and 0.018 for the reptile tree and 0.062 and 0.057 for the plant species tree, respectively (SM Table C2). As with WR in find_graphs, the 95th percentiles of *T* were higher under lineage-specific rates for both species trees. Under lineage-specific rates, these percentiles were generally higher at low ILS than at high ILS, with the highest values in the plant simulations at low ILS (Fig. 3B).

### 3.3. False detection of reticulation

Under the standard WR *≤* 3.0 criterion, find_graphs had elevated type I error rates even under constant or gene-specific substitution rates. Under a constant substitution rate, the mean type I error rate was 23.3% in the reptile simulations and 18.2% in the plant simulations. Under gene-specific rates, a type I error was made in an average of 27.9% and 17.9% replicates. Adopting the more permissive threshold WR *≤* 3.7 reduced the mean type I error rate across the two species trees to 2.7% under a constant substitution rate and to 4.2% under gene-specific rates. However, even this more permissive threshold was insufficient under lineage-specific rates, where the type I error rate averaged 80.3% for the reptile tree and 100% for the plant species tree (Fig. 3D, SM Fig. D1).

For SNaQ, the type I error rate remained low, ranging from 0% to 8% across settings. Under lineage-specific rates, SNaQ had the lowest mean type I error rate among the three methods, at 0.4% for the reptile tree and 1.3% for the plant species tree (Fig. 3D, SM Fig. D2A). The PhyNEST thresholds were calibrated using these simulations, separately for each species tree and ILS level and pooling constant and gene-specific rates. The resulting type I error rate was zero under a constant substitution rate and averaged 10% under gene-specific rates (Table D1). As with find_graphs, type I error rates remained high under lineage-specific rates despite calibration. For PhyNEST, the mean type I error rate was 86.5% in the reptile simulations and 59.0% in the plant simulations (Fig. 3D).

### 3.4. True topology recovery

Under *h* = 0, find_graphs recovered the true species tree in all reptile simulations and in nearly all plant simulations with constant or gene-specific rates. Under lineage-specific rates, recovery of the plant species tree declined to 92.5% (89%–98% across settings) (SM Fig. E1 top). Under *h* = 1, lineage-specific rates had opposite effects on topology recovery in the best-scoring graph for the two species trees. Compared with constant and gene-specific rates, the proportion of graphs displaying the true tree increased from 81.5% to 96.8% for reptiles and decreased from 76.9% to 67.8% for plants (Fig. 3C). Considering all retained graphs increased recovery for both species trees (to about 95% under constant and gene-specific rates), although recovery of the plant species tree remained lower under lineage-specific rates (73.8% versus 98.9% for reptiles; SM Fig. E1 bottom). After pruning taxon F (turtle) from the reptile phylogenies and taxon B (*Amborella*) from the plant phylogenies, the best-scoring graph displayed the corresponding pruned true tree in nearly all replicates under lineage-specific rates, but plant recovery averaged 88% to 89% under constant and gene-specific rates (SM Fig. E1 bottom right). At least one retained graph displayed the corresponding pruned true tree in at least 95% of replicates in every setting (SM Fig. E1 bottom right). The duplication/loss rate, ILS levels, and number of individuals per taxon had no noticeable effect on topology recovery (Fig. 3C).

SNaQ recovered the true topology under *h* = 0 in every replicate for both species trees. Under *h* = 1, the inferred network displayed the reptile tree less often under lineage-specific rates (82%) than under constant (87%) or gene-specific rates (96%). Recovery of the plant species tree remained high across all settings, with the inferred network displayed the true plant tree in every replicate under low ILS and in 89%–97% of replicates under high ILS (Fig. 3C). After pruning taxon F (turtle) from the reptile phylogenies and taxon B (*Amborella*) from the plant phylogenies, the networks inferred by SNaQ displayed the corresponding pruned true tree in all replicates (SM Fig. E2 top right).

The effect of lineage-specific rates on tree recovery was greater for PhyNEST than for SNaQ or find_graphs overall. Under *h* = 0, PhyNEST recovered the true topology in nearly all replicates under constant and gene-specific rates, but under lineage-specific rates, recovery fell to 35%– 66% across settings for the reptile tree and to zero for the plant species tree (SM Fig. E2 bottom). Under *h* = 1 and lineage-specific rates, the inferred network displayed the true plant species tree in only 0.5% of replicates (Fig. 3C). Under rate variation across lineages, pruning taxon F (turtle) raised recovery to 88% for reptiles, whereas recovery remained at 8% for plant after pruning taxon B (*Amborella*) (SM Fig. E2 bottom right).

### 3.5. Gene flow estimation under one hybridization event

Under *h* = 1, the minor inheritance probability *γ* estimated from the best-scoring find_graphs graph was concentrated near zero under constant and gene-specific rates. Under lineage-specific rates, the peak shifted upward to about 0.02 for the reptile simulations. The shift was much larger for the plant simulations, with the distribution peaking near 0.25. Generally, graphs that failed to display the true tree tended to have higher estimates of *γ*. This tendency was particularly visible in the plant simulations under lineage-specific rates (Fig. 4, SM Fig. F1).

**Figure 4.**
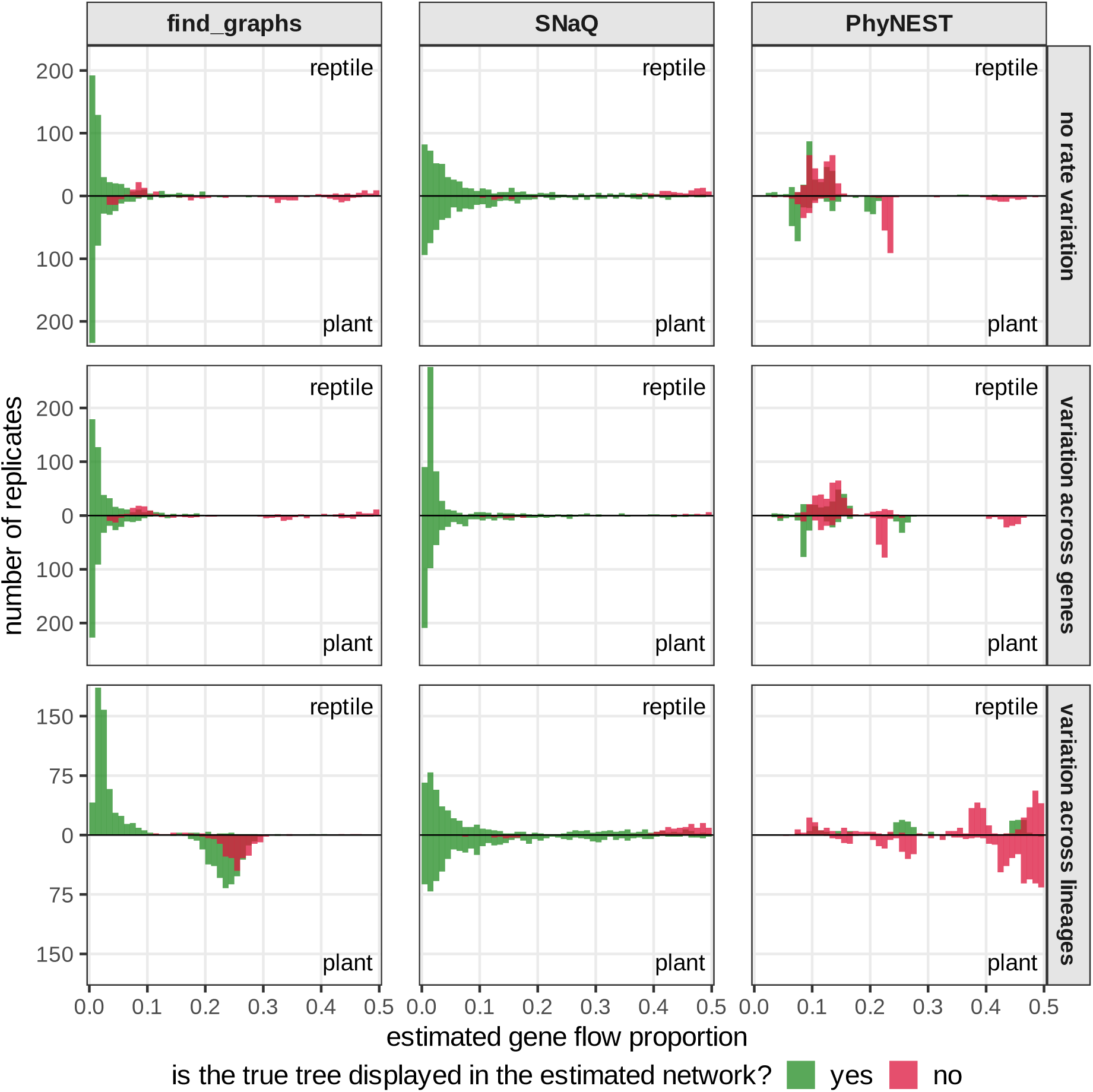
Histogram of the estimated gene flow proportion (*γ*) when allowing up to one reticulation event (*h* = 1) by find_graphs (left), SNaQ (middle) or PhyNEST (right), from data simulated under a constant rate (top), rate variation across genes (middle), or rate variation across lineages (bottom), with one individual per taxon and the three duplication/loss levels pooled. In each panel, the histogram of the reptile simulations is drawn upward and that of the plant simulations downward from the common zero horizontal line; counts are numbers of replicates (600 per species tree and panel). Histograms are colored according to whether the inferred network displays the true species tree. The two histograms are separate and their colors overlap.

For SNaQ, estimates of *γ* were concentrated near 0.02, most tightly under gene-specific rates (SM Table F1, SM Fig. F2). The upper tail (*γ* = 0.4 *−* 0.5) was more pronounced in the reptile simulations, especially under lineage-specific rates. Few networks failed to display the true tree, but of those, 90% had *γ ≥* 0.4 in the reptile simulations, and their *γ* clustered around 0.15 in the plant simulations (Fig. 4).

For PhyNEST, estimates of the minor inheritance probability *γ* were substantially farther from zero than those of find_graphs or SNaQ. Under constant and gene-specific rates, networks displaying the true plant species tree had a main peak below *γ* = 0.1 and a smaller cluster at intermediate values. Networks that failed to display the true plant species tree showed a sharper peak just above 0.2 and a tail toward 0.5. Under lineage-specific rates, the distribution of *γ* shifted toward larger values substantially, with a heavy upper tail near the maximum 0.5 (Fig. 4, SM Fig. F3). The median rose to 0.38 for the reptile simulations and 0.44 for the plant simulations (SM Table F1). Recall that this lineage-specific rate setting is when most inferred networks failed to display the true tree (section “True topology recovery”).

## 4. Discussion

Our study demonstrates that lineage-specific substitution rate variation, rather than hidden paralogy, is the primary driver of false reticulation signal in phylogenetic network inference, across our reptile and plant simulations. Under all levels of gene duplication and loss examined, including conditions in which approximately 12% of loci exhibited strong hidden paralogy involving topological discordance due to duplication and loss alone, increasing duplication and loss rates did not systematically elevate type I error rates in find_graphs, SNaQ, or PhyNEST. In contrast, lineage-specific rates frequently led find_graphs and PhyNEST to favor reticulation, although the datasets were simulated under tree topologies. For SNaQ in the reptile simulations and PhyNEST in general, networks inferred with one reticulation (*h* = 1) displayed the true species tree less often under lineage-specific rates. These findings have direct implications for the interpretation of admixture graph and phylogenetic network analyses of empirical datasets, particularly those drawn from taxa known to exhibit unequal substitution rates across lineages.

Because our simulations were calibrated from reptile UCE and plant nuclear-gene datasets, the results inform empirical animal and plant phylogenomics, using tree shapes, branch lengths, and substitution parameters estimated from real datasets. The reptile simulation parameters exhibited more extreme lineage-specific rate variation, including rapidly evolving snake and lizard lineages (Crawford et al., 2012). The plant simulation parameters had greater depth in coalescent units and represents lineages with histories of whole-genome duplication and gene loss, which can leave hidden paralogs in apparently single-copy nuclear datasets (Amborella Genome Project, 2013; Zeng et al., 2014). Accordingly, our plant simulations had more gene tree – species tree discordance due to duplications and losses. Meanwhile, consistent with the long-standing difficulty of placing turtles among amniotes (Brown and Thomson, 2017; Chiari et al., 2012; Crawford et al., 2012), taxon F in our reptile simulations (turtle) was the most difficult to resolve. Pruning it leads to near perfect topology recovery from find_graphs and SNaQ. In the plant species tree, the placement of *Amborella* is biologically important because it determines the earliest split among living plants, and it is known to be sensitive to the inference method and to the selection of the sites —slow versus fast evolving— in concatenation analyses (Xi et al., 2014). Consistent with empirical analyses, in our plant simulations the placement of taxon B (*Amborella*) was challenging, and pruning it led to an accurate recovery of the pruned species tree displayed in the estimated network.

### 4.1. Model-selection criteria require calibration

Our results show that deciding whether to add a reticulation requires a carefully calibrated threshold that is system-dependent, for both find_graphs and PhyNEST. For find_graphs, the threshold WR *≤* 3 has been applied widely across population genomics empirical studies (e.g., Ge et al., 2023; Gutaker et al., 2020; Maier et al., 2023; Zhao et al., 2023). Our results on an eight-taxon trees show that even without lineage-specific rate variation, this threshold produced substantially inflated type I error rates, leading to rejection of the tree model without reticulation. Therefore, an appropriate threshold to maintain a rate of type-1 error below 5% exceeded 3.0. Adopting a recalibrated threshold of WR *≤* 3.7, on our 8-taxon trees, controlled the type-1 error rate to remain around 5%.

For PhyNEST, we developed a model-selection procedure because the software does not provide an internal criterion for deciding whether an additional reticulation is warranted (Kong et al., 2025). We compared the per-site composite-likelihood improvement from adding a reticulation, *T*, with its distribution in simulations generated under a known tree. This reference distribution provides a cutoff for distinguishing improvements expected without reticulation from larger improvements warranting the choice of a reticulate model. The procedure offers a basis for model selection in empirical applications using PhyNEST, provided that its calibration reflects the study system.

We expect that no single threshold for either criterion is broadly appropriate. For find_graphs, the worst residual is the maximum *|z|*-score taken over all *f* -statistic residuals (comparing the observed *f* to the value expected from the graph), and the number of such residuals grows polynomially with the number of taxa. Its threshold therefore depends on the number of sampled taxa, with higher values needed to accommodate larger extremes when taking the maximum over more residuals. For PhyNEST, we accounted for alignment length by expressing the score improvement per site, but the calibrated *T^∗^* still differed between ILS levels. Thus, normalization alone does not make the decision threshold independent of the evolutionary setting. We recommend that these model-selection criteria be calibrated empirically within each study system, consistent with the recommendation for WR by Frankel and Ané (2026). For empirical applications, one possible approach is to simulate datasets under a fitted phylogeny and repeat the inference procedure to obtain the distribution of the test statistic (WR or *T*) as a null reference. This would allow calibration to reflect the sampling design and estimated evolutionary parameters. Such a simulation-based approach was developed by Cai and Ané (2021) to test how well a candidate phylogeny fits the data across 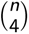 quartet CFs, based on a goodness-of-fit measure from each quartet (analogous to a *z* -score residual from each *f*_4_ statistic). This test was used for model selection on the phylogeny (tree or network) estimated with SNaQ, and led to the correct selection of a tree with high probability.

However, correct calibration of the model-selection threshold for model selection does not guarantee correct inference if the model assumptions are violated. In our simulations from the DLCoal model, gene duplications and losses did not adversely affect model selection. In contrast, rate variation across lineages increased the worst residuals in find_graphs and the score difference *T* in PhyNEST, frequently leading both criteria to favor reticulation when there was none. Calibration therefore needs to be accompanied by an assessment of the evolutionary assumptions underlying the inference. Diagnostic tests for deviations from a molecular clock (such as Tajima (1993), which evaluates a clock by testing the consistency between sequences to the outgroup using site patterns) could precede network inference to identify rate heterogeneity that could contribute to an apparent reticulation signal. Koppetsch et al. (2024) likewise recommend accounting for rate variation across lineages when testing for introgression. Their test, implemented in Dsuite (Malinsky et al., 2021), uses site patterns and spatial information to distinguish true introgression from false signal driven by rate variation: true introgression produces spatially clustered ABBA sites along chromosomes (reflecting linked variants in haplotype blocks), whereas homoplasy produces scattered sites. How to extend their robust quartet-level diagnostic test for reticulation to admixture graph inference or the composite-likelihood framework of PhyNEST is not straightforward, and would be a valuable direction for future work.

### 4.2. Lineage-specific rates drive false reticulation signal

Substitution rate variation across lineages can produce false signal of introgression through asymmetric homoplasy: when one lineage evolves faster than its sister, it independently acquires more substitutions identical to those on other branches. This creates an asymmetric excess of shared derived alleles that site-based methods cannot distinguish from genuine gene flow (Frankel and Ané, 2023; Koppetsch et al., 2024). This behavior is analogous to long-branch attraction in tree inference (Felsenstein, 1978; Roch et al., 2019), where convergent substitutions on rapidly evolving lineages mimic shared derived characters. For find_graphs, the inferred network often retained the true species tree while including a spurious admixture edge. This suggests that an additional edge can accommodate the convergent signal from the *f* -statistics without modifying the backbone tree topology. Under lineage-specific rates, PhyNEST often favored a reticulation and, unlike find_graphs, its inferred networks frequently failed to display the true species tree: its networks involved more than adding a hybrid edge to the true species tree. PhyNEST predicts site-pattern frequencies under a strict molecular clock (Kong et al., 2025). When lineages evolve at different rates, convergent substitutions can contribute to departures from these expectations. Given that PhyNEST constrains the species phylogeny to be ultrametric (unlike find_graphs), a different branching order or an added reticulation may then provide a better fit. The sensitivity of introgression methods to a violation of the molecular clock was uncovered by previous studies (Frankel and Ané, 2023, 2026; Koppetsch et al., 2024). Our results confirm this sensitivity using lineage-specific rates calibrated from empirical datasets, which were sufficient to produce severe inflation of false reticulation inference for both PhyNEST and find_graphs.

We found that ILS also influenced the response to lineage-specific rates. Under low ILS, find_graphs more frequently rejected both *h* = 0 and *h* = 1 reticulation hypotheses, whereas under high ILS a single-reticulation graph more often passed its adequacy criterion. For PhyNEST, false reticulation inference was also more frequent under low ILS. The reason why is unclear. One possibility is that under high ILS, the background noise from gene tree discordance increases the frequency of the two discordant site patterns equally, and reduces their relative difference (due to rate variation) thereby reducing the signal (interpreted as reticulation) and making a single admixture edge sufficient to absorb the residuals. Under low ILS, the rate signal may be less masked by coalescent discordance, which may drive selection of increasingly complex networks.

### 4.3. SNaQ is robust to variation in substitution rates

Across all simulation conditions, SNaQ maintained low type I error rates, including under rate variation across lineages, in line with previous reports for topology-based network methods (Cao et al., 2024; Koppetsch et al., 2024). We hypothesize that this robustness comes from SNaQ taking quartet CFs as input, which are obtained from gene estimation methods refined over decades. Fast likelihood-based methods of gene tree inference are now robust to various factors, such as a departure from a molecular clock and rate variation across loci. If gene tree topologies are unaffected by lineage-specific rates, SNaQ’s input is not affected either. Our simulations included gene tree estimation error, however, and we used these gene trees at face value to estimate quartet CFs (without integrating out gene tree uncertainty as would be possible with BUCKy, Larget et al., 2010; Solís-Lemus and Ané, 2016). Even so, the goodness-of-fit test used on the SNaQ network reliably selected the correct *h* across conditions. This robustness may come at the cost of power, however. The goodness-of-fit test on quartet CFs may be overly conservative, biased toward selecting a lower reticulation number *h* than the true value, with reduced power to detect reticulation. Indeed, Bjorner et al. (2024) found that this test rarely reject the tree-only hypothesis under deep or multiple hybridizations, as the number of taxa increased.

When allowing for *h*= 1 reticulation, the topological accuracy of SNaQ was not simply driven by the amount of gene tree estimation error, but by the type of rate variation. Gene tree estimation error was lower when rates varied across lineages, yet the true tree was displayed in the SNaQ network somewhat less often. One possible reason is that rate variation across genes leads to gene tree error that is not biased in either direction. Slow-evolving loci have low information, from too few substitutions for their tree to be reliably resolved. These uninformative loci may bias quartet CFs toward (1*/*3, 1*/*3, 1*/*3), but do not disrupt the symmetry expected from ILS alone. The asymmetry in quartet CFs, which is informative about reticulations in the network, is then driven by the well-resolved loci with a sufficiently high rate. Under lineage-specific rates, a fast-evolving lineage accumulates substitutions across all genes. If the conditions are present for a systematic bias to exist, such as long-branch attraction, then this bias affects all genes and is passed onto quartet CFs. Such bias may break the symmetry in quartet CFs expected by ILS, and a reticulation may be misleadingly favored to explain this lack of symmetry.

### 4.4. Hidden paralogy has limited impact on network inference

Our simulations included strong hidden paralogy, affecting approximately 12% of single-copy loci, that largely increased gene-species tree discordance. Yet, this topological distortion at individual loci did not systematically reduce how often inferred species phylogenies matched the true species tree or displayed it, nor did it increase false reticulation inference in our simulations. Note that we used the DLCoal model, by which gene duplications and losses follow a Poisson birth-death process along edges of the species tree. This model is most appropriate for small-scale, gene-by-gene duplication and loss events.

Previous work showed the robustness of quartet-based species tree inference methods like ASTRAL to gene duplication and loss, empirically (Aranda et al., 2026) and theoretically under a gene duplication and loss model without ILS (Legried et al., 2021) and with ILS (Hill et al., 2022; Markin and Eulenstein, 2021). These theoretical works established that the gene duplication/loss models they considered maintain the symmetry of quartet CFs when the species phylogeny is a tree, by quantifying the frequency of strong hidden paralogy. SNaQ uses the same quartet CFs (topology frequencies) as ASTRAL, so its robustness to the presence of strong hidden paralogy is not surprising, at least under a tree model (*h* = 0). The robustness of methods using quartet CFs may continue to hold when the species phylogeny is a network with reticulations if duplications and losses do not disrupt the symmetries expected from ILS on the network, but theory beyond the tree case is lacking.

However, we expect that our conclusions are dependent on the model we chose to simulate duplications and losses (DLCoal), applied uniformly across the species tree, with constant rates of gene duplications and losses across lineages. We did not simulate whole-genome duplication (WGD) or large-scale segmental duplication events, for example. After a WGD, gene loss is particularly elevated, in a process called gene fractionation (Langham et al., 2004). Gene loss is not independent across copies from the two paralog loci: once one copy is lost, the other copy tends to be retained since losing both copies is typically deleterious due to selective pressure. Also, gene loss from blocks of duplicated genes is often non-uniform following a WGD, as physically interacting genes tend to be co-retained (Makino and McLysaght, 2012). If all lost copies are from the same paralog (e.g. all from the duplicated locus), then orthology is preserved. If copies are lost from both paralogs instead, then strong hidden paralogy can be frequent. If speciation occurs shortly after a WGD and before gene fractionation is mostly complete, then gene loss has the opportunity to vary across lineages, and strong hidden paralogy might be elevated. Xiong et al. (2022) showed that strong hidden paralogy can be particularly elevated if the loss rate varies across lineages differentially across the two paralog loci, that is, if the gene copy from one paralog has a high loss rate in one lineage while the copy from the other paralog has a high loss rate in other lineages. Under the tractable WGD model of Rabier et al. (2014), fractionation occurs immediately after the WGD and gene loss is assumed independent across gene families, such that the symmetry of quartet CFs expected under ILS alone may be preserved. However, as mentioned above, there is empirical evidence that gene losses can be non-independent, fractionation may span speciation events, and the rate of gene loss may vary across lineages. Although the plant species tree includes lineages with histories of WGD, duplications and losses were simulated under the same DLCoal process for both species trees. The similar outcomes therefore support robustness to small-scale duplication and loss under DLCoal in both sets of simulations, but do not directly establish robustness to WGD and subsequent fractionation. Theoretical work is needed under a realistic WGD model with variable rates of gene duplication/loss across lineages and under reticulate species networks, to determine the expected behavior and symmetry of quartet CFs, and whether species phylogeny inference methods remain consistent.

### 4.5. Conclusion

This study evaluated two model violations that have been proposed as potential sources of false reticulation signal: hidden paralogy from gene duplication and loss, and substitution rate variation across lineages in simulations designed from reptile and plant datasets. Hidden paralogy did not mislead network inference under the conditions we tested. Lineage-specific rate variation, in contrast, misled all three methods in different ways.

SNaQ maintained near-perfect type-I error but recovered the true topology less often when lineages had variable rates. From *f* -statistics, find_graphs systematically selected spurious reticulations. Even under constant rates, elevated worst residuals can indicate rate-induced artifacts rather than genuine reticulation. For PhyNEST, our calibrated score-difference criterion provides a basis for choosing between tree and single-reticulation models For both methods, we recommend calibrating the threshold to the study system, as this helped limit false positives under constant and gene-specific rates. However, we still recommend caution because false reticulations remained frequent when the molecular clock was violated.

Finally, when a reticulation was allowed, a high estimated gene flow proportion (*γ* approaching 0.5) could accompany networks that failed to display the true species tree. The implication for empirical analyses is that a large admixture (or gene flow) proportion should be interpreted with caution: it may either signal a true hybridization or admixture event with an overwhelming effect on half of the genome. Alternatively, it may be a rate-induced artifact, with an extra erroneous reticulation event, on a distorted network that does not display the true species topology when the gene flow edge with large *γ* is removed. Therefore, we recommend assessing the validity of a molecular clock in the system under study, and interpreting the results carefully if rate heterogeneity across lineages is plausible.

## Acknowledgements

We thank Max Sherwin for his early contributions to this project, including initial coding efforts that helped guide its development. We also thank Christopher Blair, Nicole Haderlein and Ayesha Omar for discussions about reptile and turtle evolution, and prior joint work on the Crawford et al., 2012 data. We sincerely appreciate Sarah Friehrich for graphic support.

## Data, script, and code availability

The code to reproduce all analyses and figures is available on GitHub at https://github.com/ cecileane/simulation-hiddenparalogy-ratevariation and in Dryad (doi: 10.5061/dryad.dfn2z35h7).

## Supplementary Material A. Simulation workflow

Figure A1 provides technical details on the gene tree simulation that were omitted from the Methods. For each replicate, SimPhy was run iteratively in batches. After each batch, gene trees with repeated or missing taxa were discarded. Valid trees were accumulated across batches until *K* valid trees where simulated with *K ≥* 1000, or the iteration limit of 20,000 was reached. If fewer than 800 valid trees were obtained, the replicate was considered failed. No replicate failed under any parameter setting in this study. At least min(1000, *K*) valid trees were passed to Seq-Gen.

**Figure A1.**
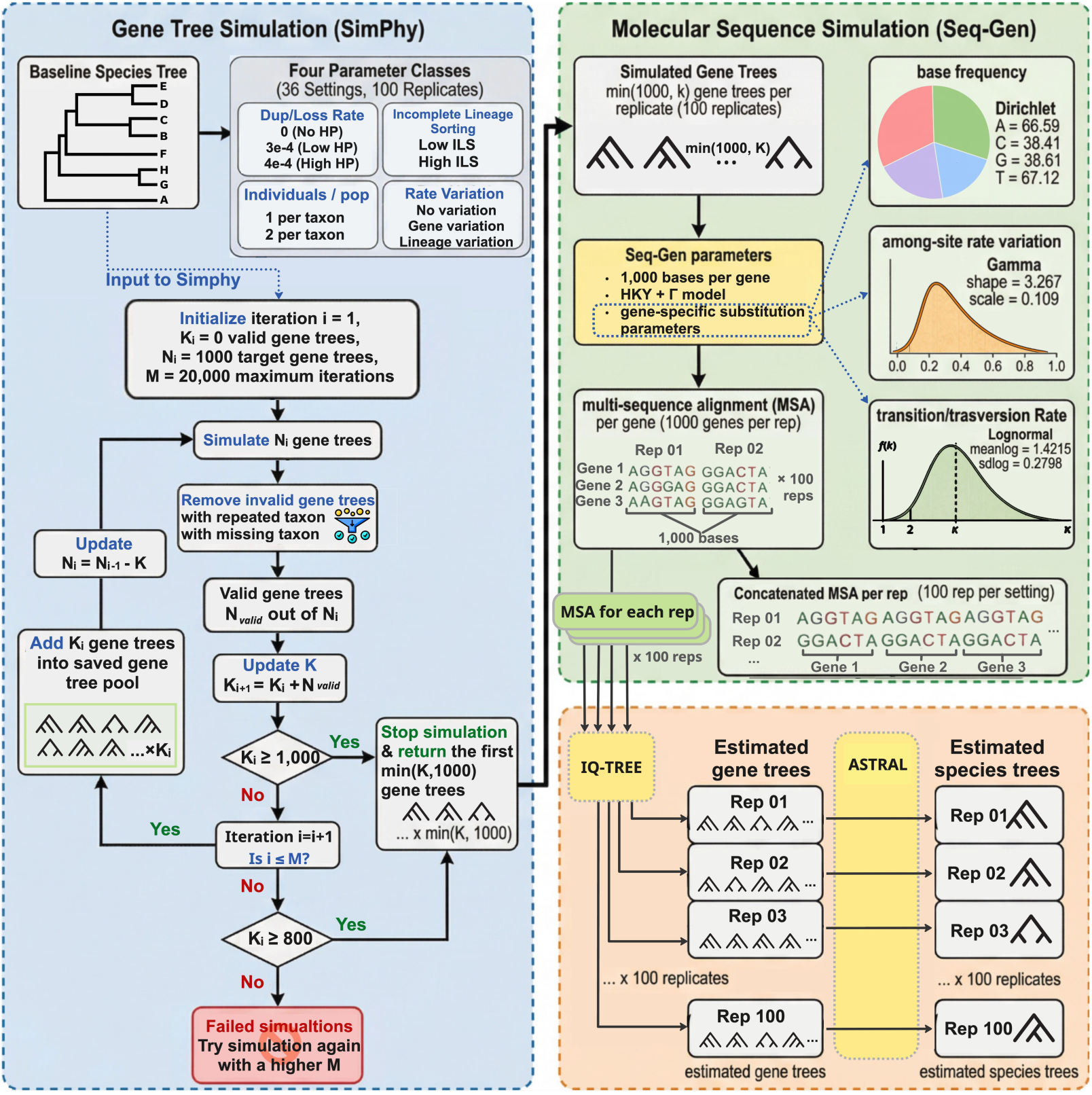
Overview of the full pipeline: gene tree simulation (SimPhy), sequence simulation (Seq-Gen), gene tree estimation (IQ-TREE), and species tree estimation (ASTRAL)

## Supplementary Material B. Gene tree variability

**Figure B1.**
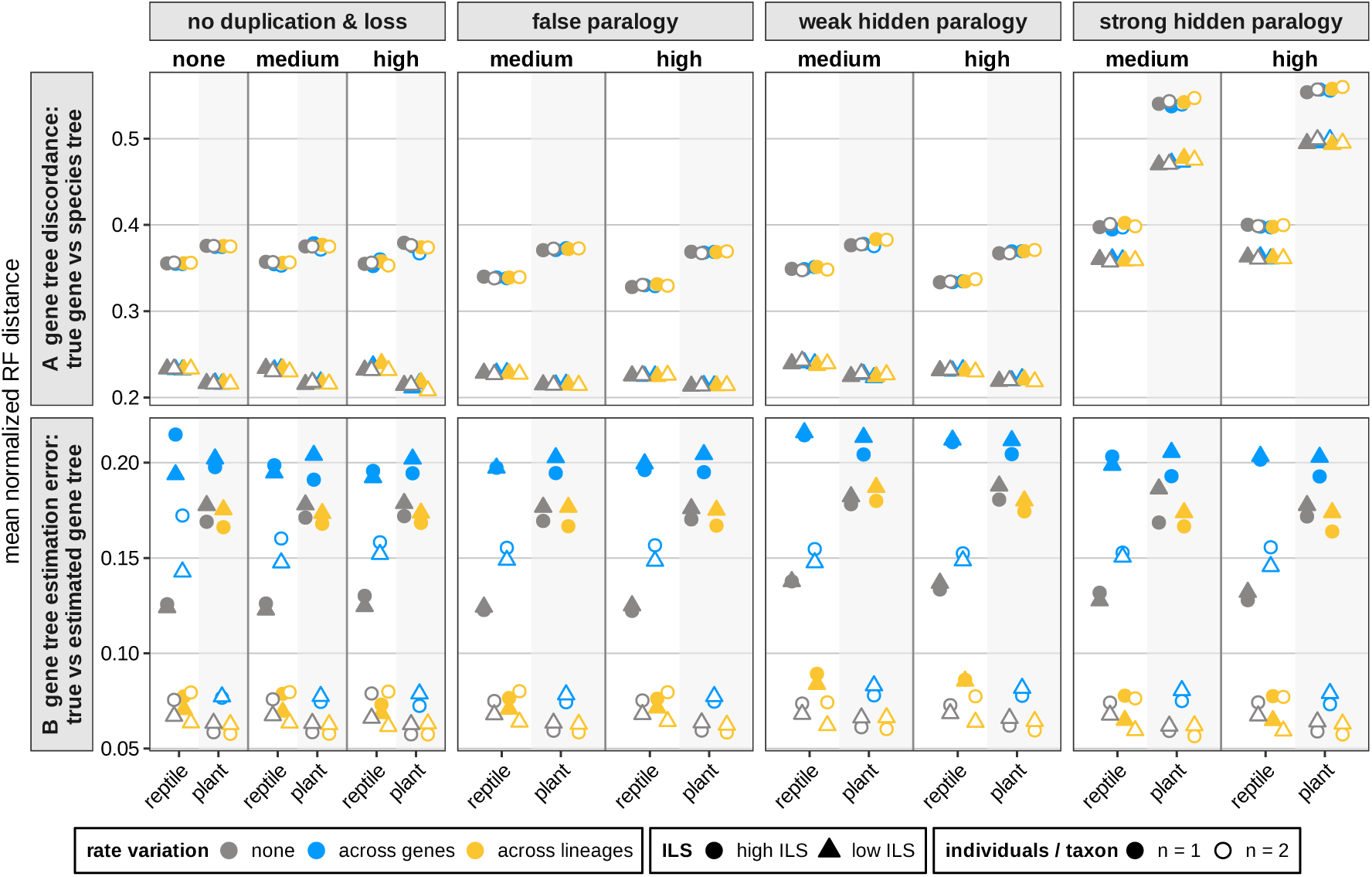
Gene tree discordance and estimation error. Top: average normalized Robinson–Foulds (RF) distance between the true gene tree and the true species tree, measuring gene tree discordance. Bottom: average normalized RF distance between the true gene tree and the gene tree estimated by IQ-TREE, measuring gene tree estimation error. The x-axis separates the reptile and the plant species tree within each duplication/loss level (none, medium, high: 0, 3 *×* 10*^−^*^4^ and 4 *×* 10*^−^*^4^ for the reptile tree, 0, 1 *×* 10*^−^*^4^ and 1.4 *×* 10*^−^*^4^ for the plant species tree). Columns separate loci of different type: orthologous or with hidden paralogy (which requires a non-zero duplication and loss rate). Color: substitution rate variation; shape: ILS level; filled and open symbols: one and two individuals per taxon.

**Table B1.** Species tree – gene tree discordance: normalized RF distance between the true (SimPhy) gene tree and the species tree. All distances are between 8-taxon trees. With one individual per taxon, the gene tree was compared directly to the species tree. With two individuals per taxon, the two individuals of a taxon were first merged into a single tip when they were monophyletic in the gene tree, to decrease computing time. Then, all subtrees obtained by sampling 1 individual per taxon (up to 2^8^ subtrees) were compared to the species tree, and their RF distances to the species tree were averaged. Each cell combines the 1800 or 1200 replicates in the corresponding simulation setting (from 3 rate variation categories, 1 or 2 individuals/taxon, and either 3 or 2 duplication rates as paralogs never occurred under a rate of 0). The last four column pairs give the percentage of gene trees with exactly 0 (identical to the species tree), 1, 2 or at least 3 discordant branches. With 2 individuals per taxon, this is the percentage of subtrees sampling one individual per taxon, with equal weight to all 2^8^ subtrees per gene tree.

|  |  | mean normalized RF |  | % of gene trees by number of discordant branches |  |  |  |  |  |  |  |
|  |  |  |  | 0 (identical) |  | 1 |  | 2 |  | ≥ 3 |  |
|---|---|---|---|---|---|---|---|---|---|---|---|
| locus category | ILS | reptile | plant | reptile | plant | reptile | plant | reptile | plant | reptile | plant |
| orthologs | high | 0.356 | 0.375 | 9.6 | 10.2 | 28.0 | 27.9 | 40.8 | 33.4 | 21.7 | 28.5 |
|  | low | 0.233 | 0.216 | 22.5 | 27.9 | 41.4 | 43.0 | 33.4 | 22.9 | 2.8 | 6.1 |
| false paralogs | high | 0.334 | 0.370 | 11.4 | 10.5 | 30.0 | 28.6 | 40.8 | 33.2 | 17.8 | 27.7 |
|  | low | 0.227 | 0.214 | 23.7 | 28.0 | 41.5 | 43.4 | 32.6 | 22.6 | 2.2 | 6.0 |
| weak hidden paralogs | high | 0.342 | 0.374 | 13.0 | 10.8 | 29.0 | 28.0 | 35.7 | 32.8 | 22.3 | 28.4 |
|  | low | 0.236 | 0.223 | 24.5 | 27.6 | 39.4 | 41.4 | 30.0 | 23.8 | 6.0 | 7.2 |
| strong hidden paralogs | high | 0.399 | 0.549 | 2.8 | 0.9 | 23.5 | 11.6 | 48.8 | 27.6 | 24.9 | 59.9 |
|  | low | 0.361 | 0.485 | 3.1 | 1.3 | 30.2 | 20.0 | 51.1 | 33.1 | 15.6 | 45.6 |

**Table B2.**
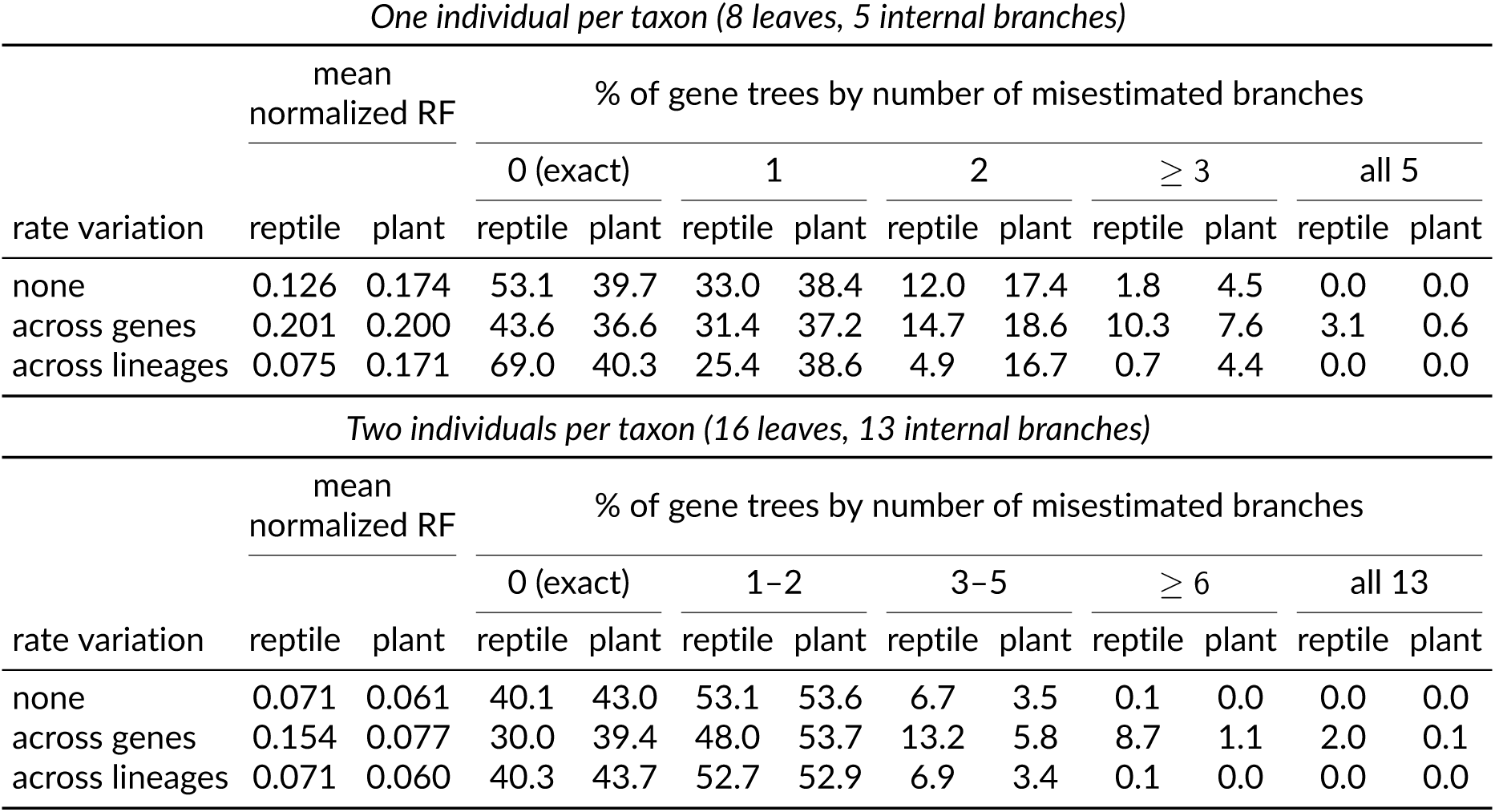
Gene tree estimation error: normalized RF distance between the true (SimPhy) gene tree and the gene tree estimated by IQ-TREE. Each cell summarizes 600 replicate datasets (from 3 duplication/loss rates *×* 2 ILS levels). The normalized RF distance is the raw distance divided by its maximum 2(8*n*_ind_ *−* 3): 5 branches with *n*_ind_ = 1 individual per taxon, 13 with *n*_ind_ = 2. The remaining column pairs give the percentage of gene trees by number of misestimated internal branches; the tiers differ between the two panels because the trees have 5 and 13 internal branches respectively.

| *One individual per taxon (8 leaves, 5 internal branches)* |  |  |  |  |  |  |  |  |  |  |  |
|  | mean normalized RF |  | % of gene trees by number of misestimated branches |  |  |  |  |  |  |  |  |
|  |  |  | 0 (exact) |  | 1 |  | 2 |  | ≥ 3 |  | all 5 |  |
| rate variation | reptile | plant | reptile | plant | reptile | plant | reptile | plant | reptile | plant | reptile | plant |
|---|---|---|---|---|---|---|---|---|---|---|---|---|
| none | 0.126 | 0.174 | 53.1 | 39.7 | 33.0 | 38.4 | 12.0 | 17.4 | 1.8 | 4.5 | 0.0 | 0.0 |
| across genes | 0.201 | 0.200 | 43.6 | 36.6 | 31.4 | 37.2 | 14.7 | 18.6 | 10.3 | 7.6 | 3.1 | 0.6 |
| across lineages | 0.075 | 0.171 | 69.0 | 40.3 | 25.4 | 38.6 | 4.9 | 16.7 | 0.7 | 4.4 | 0.0 | 0.0 |
| *Two individuals per taxon (16 leaves, 13 internal branches)* |  |  |  |  |  |  |  |  |  |  |  |  |
|  | mean normalized RF |  | % of gene trees by number of misestimated branches |  |  |  |  |  |  |  |  |  |
|  |  |  | 0 (exact) |  | 1–2 |  | 3–5 |  | ≥ 6 |  | all 13 |  |
| rate variation | reptile | plant | reptile | plant | reptile | plant | reptile | plant | reptile | plant | reptile | plant |
| none | 0.071 | 0.061 | 40.1 | 43.0 | 53.1 | 53.6 | 6.7 | 3.5 | 0.1 | 0.0 | 0.0 | 0.0 |
| across genes | 0.154 | 0.077 | 30.0 | 39.4 | 48.0 | 53.7 | 13.2 | 5.8 | 8.7 | 1.1 | 2.0 | 0.1 |
| across lineages | 0.071 | 0.060 | 40.3 | 43.7 | 52.7 | 52.9 | 6.9 | 3.4 | 0.1 | 0.0 | 0.0 | 0.0 |

## Supplementary Material C. Distribution of the test statistic for model selection

This section presents supplementary summaries of the empirical distribution of the worst residual (WR) from find_graphs and the per-site score difference *T* from PhyNEST, which are the test statistics used for model selection.

**Figure C1.**
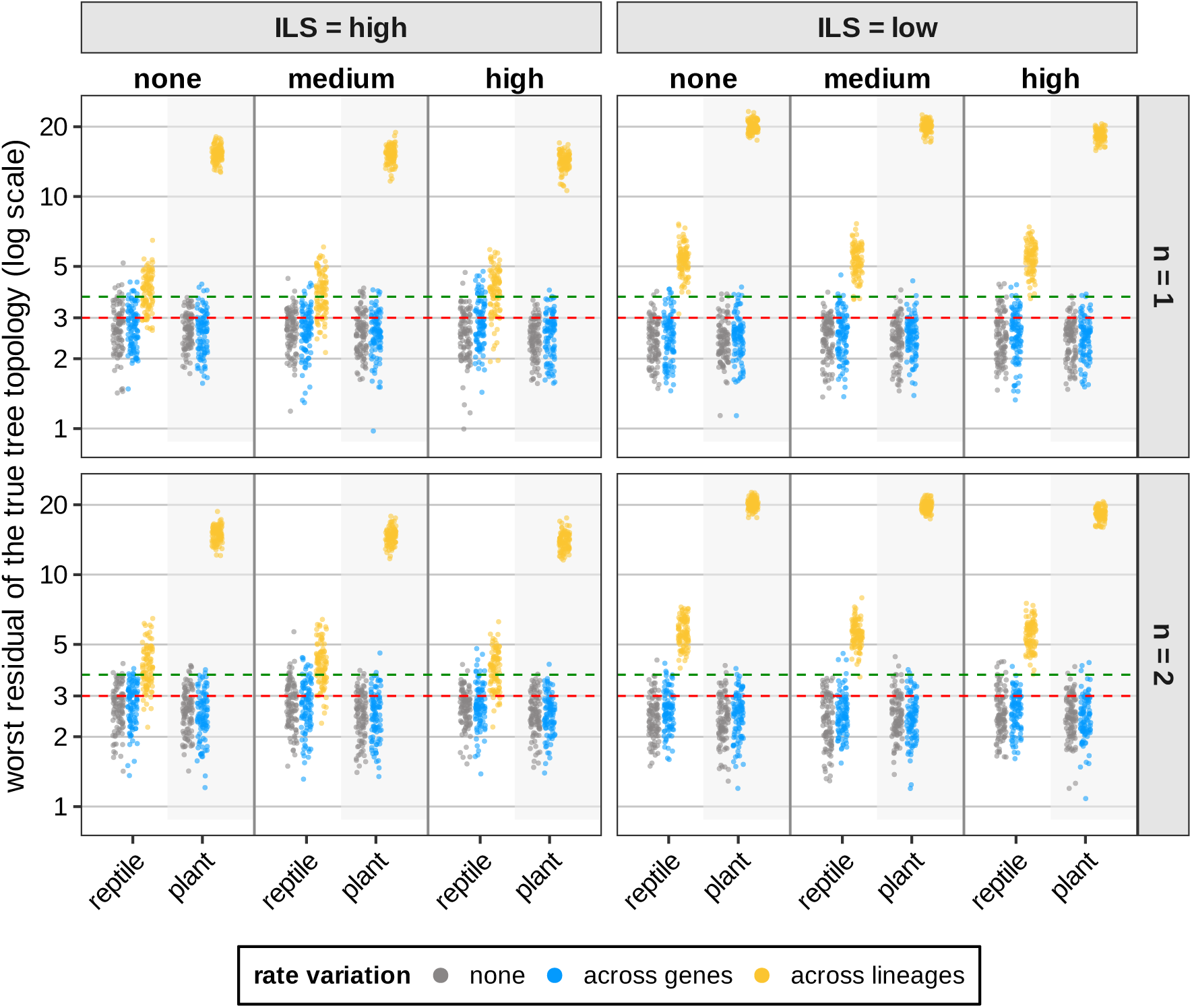
Worst residual (WR) of the true species tree topology, with branch lengths fitted to the *f* -statistics, in every replicate analyzed with find_graphs (log scale). Columns separate incomplete lineage sorting (ILS) levels and rows the number of individuals sampled per taxon (*n*). Within each panel, the x-axis separates the duplication/loss level (none, medium, high: 0, 3*×*10*^−^*^4^ and 4*×*10*^−^*^4^ for the reptile tree, 0, 1*×*10*^−^*^4^ and 1.4*×*10*^−^*^4^ for the plant species tree), with the reptile and the plant species tree side by side. Colors indicate substitution rate variation scenarios: across genes (blue), across lineages (yellow), and none (gray). Dashed horizontal lines mark the thresholds of worst residual (WR = 3.0 in red and WR = 3.7 in green) used here for model selection.

**Figure C2.**
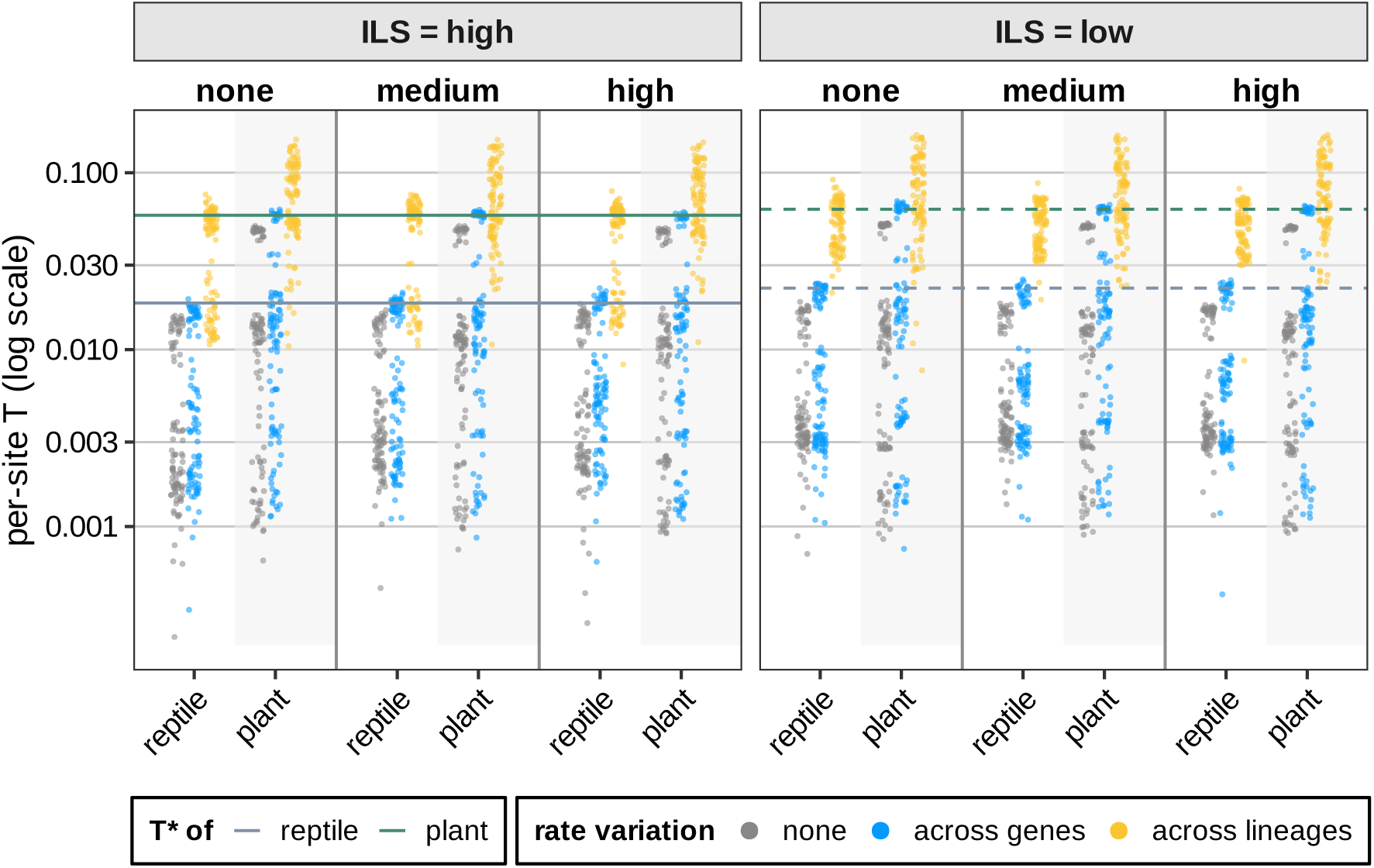
Per-site score difference *T* from PhyNEST in every replicate (one individual per taxon; log scale), where *T* = (score*_h_*_=0_ *−* score*_h_*_=1_)*/M* and *M* is the number of sites in the concatenated alignment. Columns separate ILS levels. Within each panel, the x-axis separates the duplication/loss level (none, medium, high), with the reptile and the plant species tree side by side. Colors indicate substitution rate variation scenarios as in Fig. C1. Horizontal lines mark the calibrated threshold *T^∗^* used to select *h* = 1 when *T > T^∗^* (dashed: reptile, 0.018 under high and 0.022 under low ILS; solid: plant, 0.057 under high and 0.062 under low ILS).

**Table C1.**
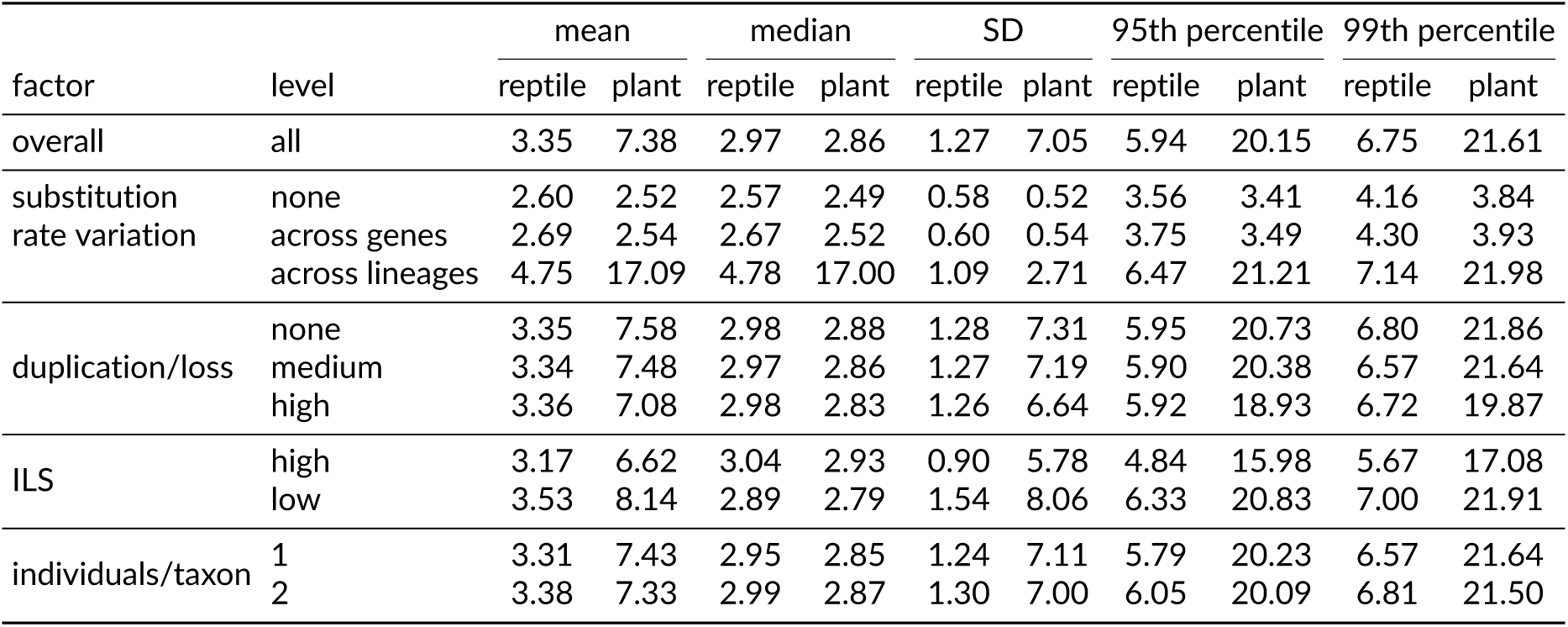
Summary statistics of the worst residual (WR) of the true species tree from find_graphs. Statistics (mean, median, standard deviation, 95th and 99th percentiles) are computed over all replicate WR values pooled within each factor level (100 replicates per setting, 36 settings per tree).

| factor | level | mean |  | median |  | SD |  | 95th percentile |  | 99th percentile |  |
|---|---|---|---|---|---|---|---|---|---|---|---|
|  |  | reptile | plant | reptile | plant | reptile | plant | reptile | plant | reptile | plant |
| overall | all | 3.35 | 7.38 | 2.97 | 2.86 | 1.27 | 7.05 | 5.94 | 20.15 | 6.75 | 21.61 |
| substitution rate variation | none | 2.60 | 2.52 | 2.57 | 2.49 | 0.58 | 0.52 | 3.56 | 3.41 | 4.16 | 3.84 |
|  | across genes | 2.69 | 2.54 | 2.67 | 2.52 | 0.60 | 0.54 | 3.75 | 3.49 | 4.30 | 3.93 |
|  | across lineages | 4.75 | 17.09 | 4.78 | 17.00 | 1.09 | 2.71 | 6.47 | 21.21 | 7.14 | 21.98 |
| duplication/loss | none | 3.35 | 7.58 | 2.98 | 2.88 | 1.28 | 7.31 | 5.95 | 20.73 | 6.80 | 21.86 |
|  | medium | 3.34 | 7.48 | 2.97 | 2.86 | 1.27 | 7.19 | 5.90 | 20.38 | 6.57 | 21.64 |
|  | high | 3.36 | 7.08 | 2.98 | 2.83 | 1.26 | 6.64 | 5.92 | 18.93 | 6.72 | 19.87 |
| ILS | high | 3.17 | 6.62 | 3.04 | 2.93 | 0.90 | 5.78 | 4.84 | 15.98 | 5.67 | 17.08 |
|  | low | 3.53 | 8.14 | 2.89 | 2.79 | 1.54 | 8.06 | 6.33 | 20.83 | 7.00 | 21.91 |
| individuals/taxon | 1 | 3.31 | 7.43 | 2.95 | 2.85 | 1.24 | 7.11 | 5.79 | 20.23 | 6.57 | 21.64 |
|  | 2 | 3.38 | 7.33 | 2.99 | 2.87 | 1.30 | 7.00 | 6.05 | 20.09 | 6.81 | 21.50 |

**Table C2.**
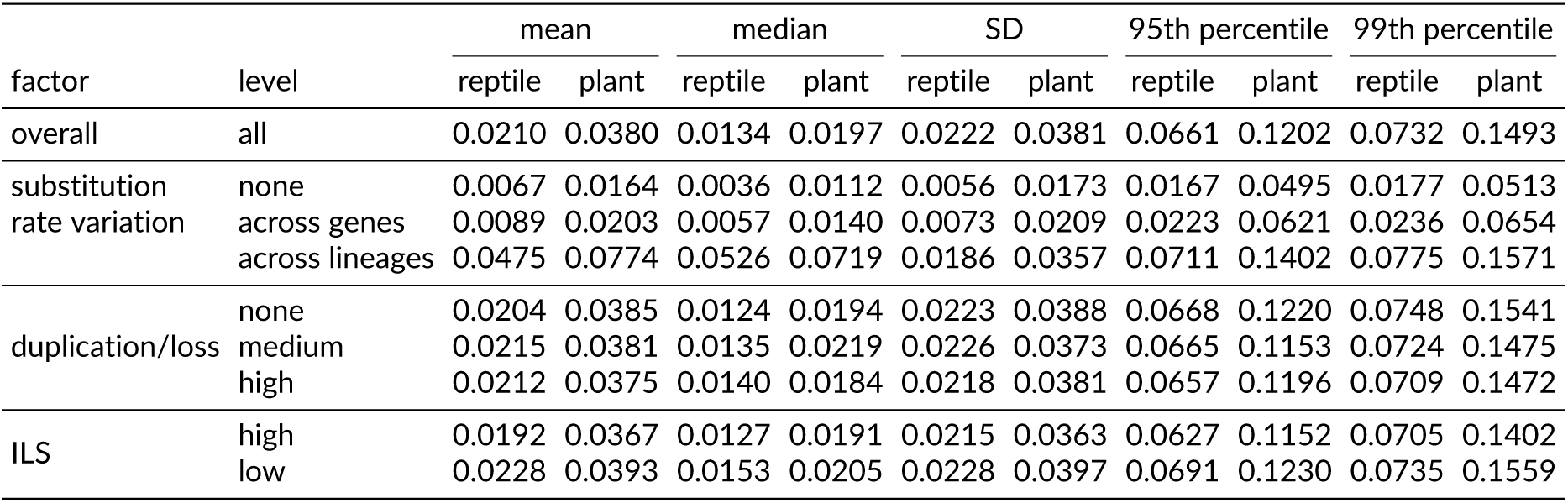
Summary statistics of the per-site score difference *T* from PhyNEST. Statistics (mean, median, standard deviation, 95th and 99th percentiles) are computed over all replicate values of *T* pooled within each factor level. The calibrated thresholds *T^∗^*, the 95th percentiles of *T* pooled over the settings without lineage-specific rates, are 0.018 under high and 0.022 under low ILS for the reptile tree, and 0.057 under high and 0.062 under low ILS for the plant species tree.

| factor | level | mean |  | median |  | SD |  | 95th percentile |  | 99th percentile |  |
|---|---|---|---|---|---|---|---|---|---|---|---|
|  |  | reptile | plant | reptile | plant | reptile | plant | reptile | plant | reptile | plant |
| overall | all | 0.0210 | 0.0380 | 0.0134 | 0.0197 | 0.0222 | 0.0381 | 0.0661 | 0.1202 | 0.0732 | 0.1493 |
| substitution rate variation | none | 0.0067 | 0.0164 | 0.0036 | 0.0112 | 0.0056 | 0.0173 | 0.0167 | 0.0495 | 0.0177 | 0.0513 |
|  | across genes | 0.0089 | 0.0203 | 0.0057 | 0.0140 | 0.0073 | 0.0209 | 0.0223 | 0.0621 | 0.0236 | 0.0654 |
|  | across lineages | 0.0475 | 0.0774 | 0.0526 | 0.0719 | 0.0186 | 0.0357 | 0.0711 | 0.1402 | 0.0775 | 0.1571 |
| duplication/loss | none | 0.0204 | 0.0385 | 0.0124 | 0.0194 | 0.0223 | 0.0388 | 0.0668 | 0.1220 | 0.0748 | 0.1541 |
|  | medium | 0.0215 | 0.0381 | 0.0135 | 0.0219 | 0.0226 | 0.0373 | 0.0665 | 0.1153 | 0.0724 | 0.1475 |
|  | high | 0.0212 | 0.0375 | 0.0140 | 0.0184 | 0.0218 | 0.0381 | 0.0657 | 0.1196 | 0.0709 | 0.1472 |
| ILS | high | 0.0192 | 0.0367 | 0.0127 | 0.0191 | 0.0215 | 0.0363 | 0.0627 | 0.1152 | 0.0705 | 0.1402 |
|  | low | 0.0228 | 0.0393 | 0.0153 | 0.0205 | 0.0228 | 0.0397 | 0.0691 | 0.1230 | 0.0735 | 0.1559 |

## Supplementary Material D. Model selection

**Figure D1.**
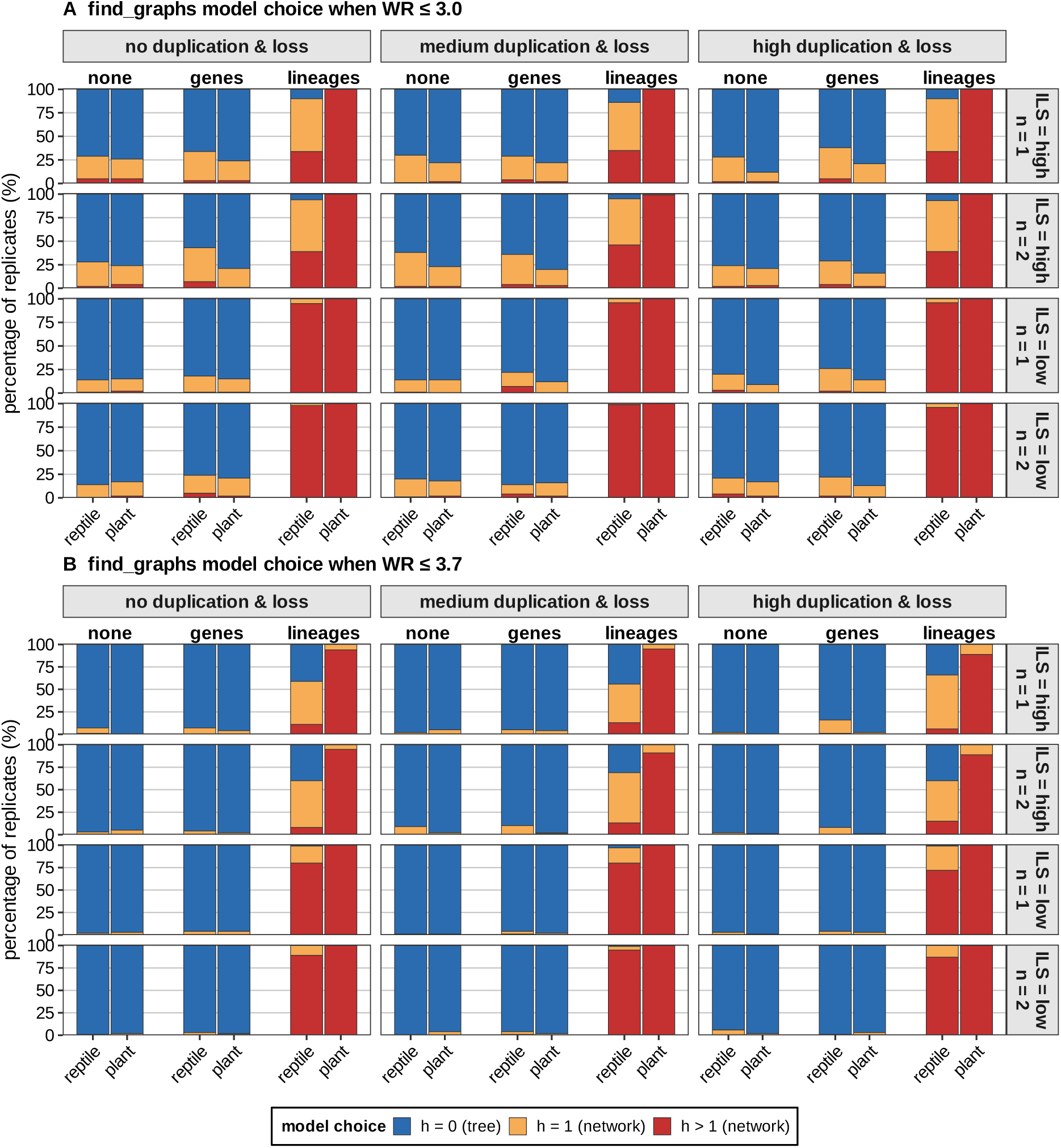
Model choice by find_graphs under two worst residual thresholds (A: WR *≤* 3.0; B: WR *≤* 3.7), for the reptile tree (left bar of each pair) and the plant species tree (right bar). Each bar shows the percentage of replicates selecting each model, with colors indicating the inferred number of hybridization events: *h* = 0 (tree, blue), *h* = 1 (orange), and *h >* 1 (red). The x-axis separates different substitution rate variation (none, gene-specific, or lineage-specific). Columns show the duplication/loss level (none, medium, high). Rows show the ILS level (high or low) and the number of individuals sampled per taxon (1 or 2).

**Figure D2.**
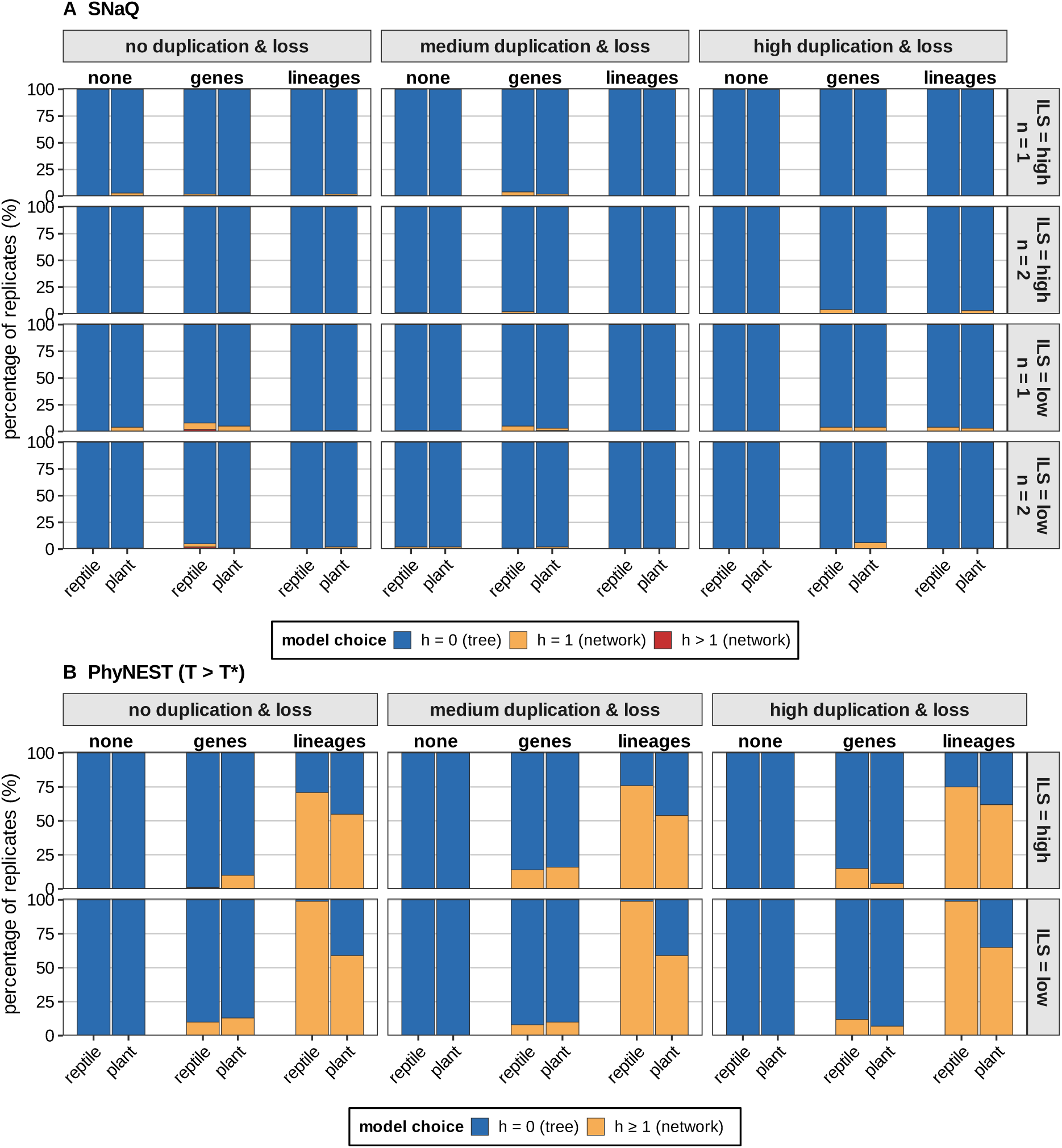
Model choice by (A) SNaQ and (B) PhyNEST under the calibrated rule (*h ≥* 1 if *T > T^∗^*, with *T^∗^* = 0.022 and 0.018 under low and high ILS for the reptile tree, 0.062 and 0.057 for the plant species tree), for the reptile tree (left bar of each pair) and the plant species tree (right bar). Each bar shows the percentage of replicates selecting each model, with colors indicating the inferred number of hybridization events: *h* = 0 (tree, blue), *h* = 1 (orange), and *h >* 1 (red) for SNaQ; for PhyNEST, orange denotes rejecting *h* = 0, that is, favoring *h ≥* 1. The x-axis separates different substitution rate variation (none, gene-specific, or lineage-specific). Columns show the duplication/loss level (none, medium, high). Rows show the ILS level (high or low) and, in (A), the number of individuals sampled per taxon (1 or 2).

**Table D1.** Percentage of replicates favoring *h* = 1 and *h >* 1. The sum of the *h* = 1 and *h >* 1 columns gives the overall rate of type I error (favoring reticulation *h ≥* 1 instead of a tree with *h* = 0). Results are compared across three methods: find_graphs applied under two worst residual thresholds (WR *≤* 3.0 and WR *≤* 3.7), SNaQ, and PhyNEST under the calibrated rule *T > T^∗^* (one individual per taxon; a rejection of *h* = 0 is reported as favoring *h ≥* 1). Within each method, percentages are across all simulation replicates sharing the indicated parameter level.

| find_graphs |  |  |  |  |  |  |  |  |  |
|---|---|---|---|---|---|---|---|---|---|
|  |  | WR ≤ 3.0, *h* = 1 |  | WR ≤ 3.0, *h* > 1 |  | WR ≤ 3.7, *h* = 1 |  | WR ≤ 3.7, *h* > 1 |  |
| factor | level | reptile | plant | reptile | plant | reptile | plant | reptile | plant |
| overall | all | 24.6 | 10.8 | 24.4 | 34.5 | 13.9 | 2.8 | 15.9 | 32.1 |
| rate variation | none | 21.3 | 16.0 | 2.0 | 2.2 | 3.1 | 2.2 | 0.1 | 0.0 |
|  | across genes | 23.9 | 16.4 | 4.0 | 1.5 | 5.6 | 2.4 | 0.2 | 0.2 |
|  | across lineages | 28.4 | 0.1 | 67.2 | 99.9 | 32.9 | 3.9 | 47.4 | 96.1 |
| duplication/loss | none | 24.8 | 11.9 | 24.2 | 35.0 | 13.3 | 2.7 | 15.8 | 32.5 |
|  | medium | 23.7 | 11.2 | 25.0 | 34.3 | 12.8 | 2.9 | 16.9 | 32.2 |
|  | high | 25.2 | 9.3 | 24.1 | 34.2 | 15.5 | 2.9 | 15.1 | 31.5 |
| ILS | high | 37.0 | 12.4 | 14.9 | 34.9 | 20.9 | 4.1 | 3.8 | 30.8 |
|  | low | 12.1 | 9.2 | 33.9 | 34.2 | 6.8 | 1.6 | 28.1 | 33.4 |
| individuals/taxon | 1 | 24.6 | 10.4 | 23.6 | 34.3 | 14.9 | 2.8 | 14.7 | 32.1 |
|  | 2 | 24.5 | 11.2 | 25.2 | 34.7 | 12.8 | 2.8 | 17.1 | 32.1 |
| SNaQ **and** PhyNEST |  |  |  |  |  |  |  |  |  |
|  |  | SNaQ *h* = 1 |  | SNaQ *h* > 1 |  | PhyNEST *h* ≥ 1 |  |  |  |
| factor | level | reptile | plant | reptile | plant | reptile | plant |  |  |
| overall | all | 1.2 | 1.4 | 0.1 | 0.1 | 32.2 | 23.0 |  |  |
| rate variation | none | 0.5 | 1.2 | 0.0 | 0.0 | 0.0 | 0.0 |  |  |
|  | across genes | 2.6 | 2.0 | 0.3 | 0.1 | 10.0 | 10.0 |  |  |
|  | across lineages | 0.4 | 1.2 | 0.0 | 0.1 | 86.5 | 59.0 |  |  |
| duplication/loss | none | 1.0 | 1.8 | 0.3 | 0.1 | 30.2 | 22.8 |  |  |
|  | medium | 1.3 | 1.0 | 0.0 | 0.1 | 32.8 | 23.2 |  |  |
|  | high | 1.2 | 1.6 | 0.0 | 0.0 | 33.5 | 23.0 |  |  |
| ILS | high | 0.8 | 0.8 | 0.0 | 0.1 | 28.0 | 22.3 |  |  |
|  | low | 1.5 | 2.1 | 0.2 | 0.1 | 36.3 | 23.7 |  |  |
| individuals/taxon | 1 | 1.6 | 1.7 | 0.1 | 0.1 | 32.2 | 23.0 |  |  |
|  | 2 | 0.8 | 1.2 | 0.1 | 0.0 | – | – |  |  |

## Supplementary Material E. Recovery of the true tree as displayed in the inferred

**Figure E1.**
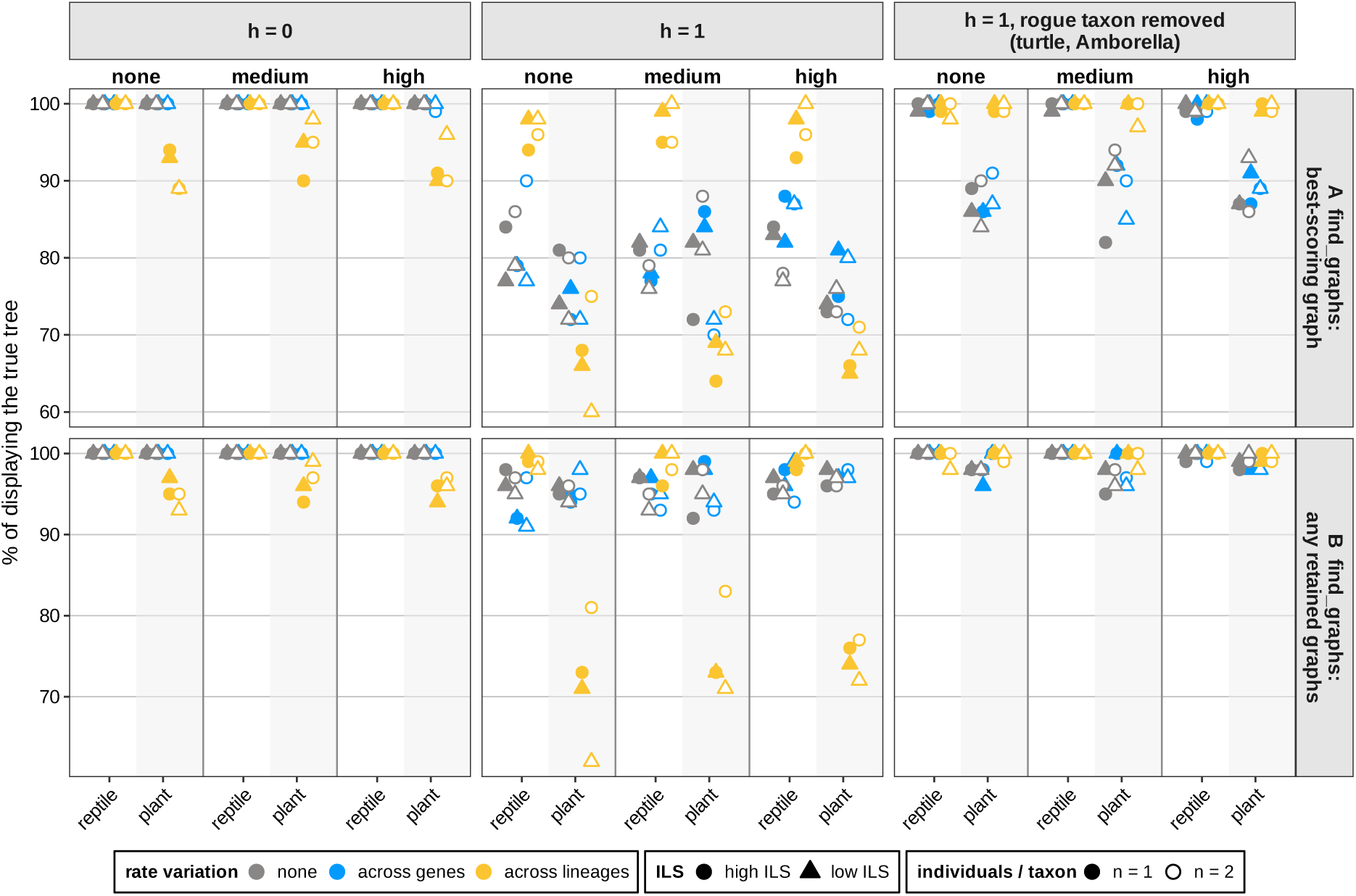
Probability that the true species topology is displayed in (A) the best-scoring graph from find_graphs, and (B) one of the retained graphs from find_graphs. Note that the y-axis has different scales across panels. Columns correspond to *h* = 0, *h* = 1, and *h* = 1 with the rogue taxon pruned from the true and estimated networks: taxon F (turtle) for the reptile tree and taxon B (*Amborella*) for the plant species tree. Duplication/loss rates (none, medium, high) are in different subpanels, colors indicate substitution rate variation, shapes the level of ILS, and filled versus open symbols the number of individuals per taxon.

**Figure E2.**
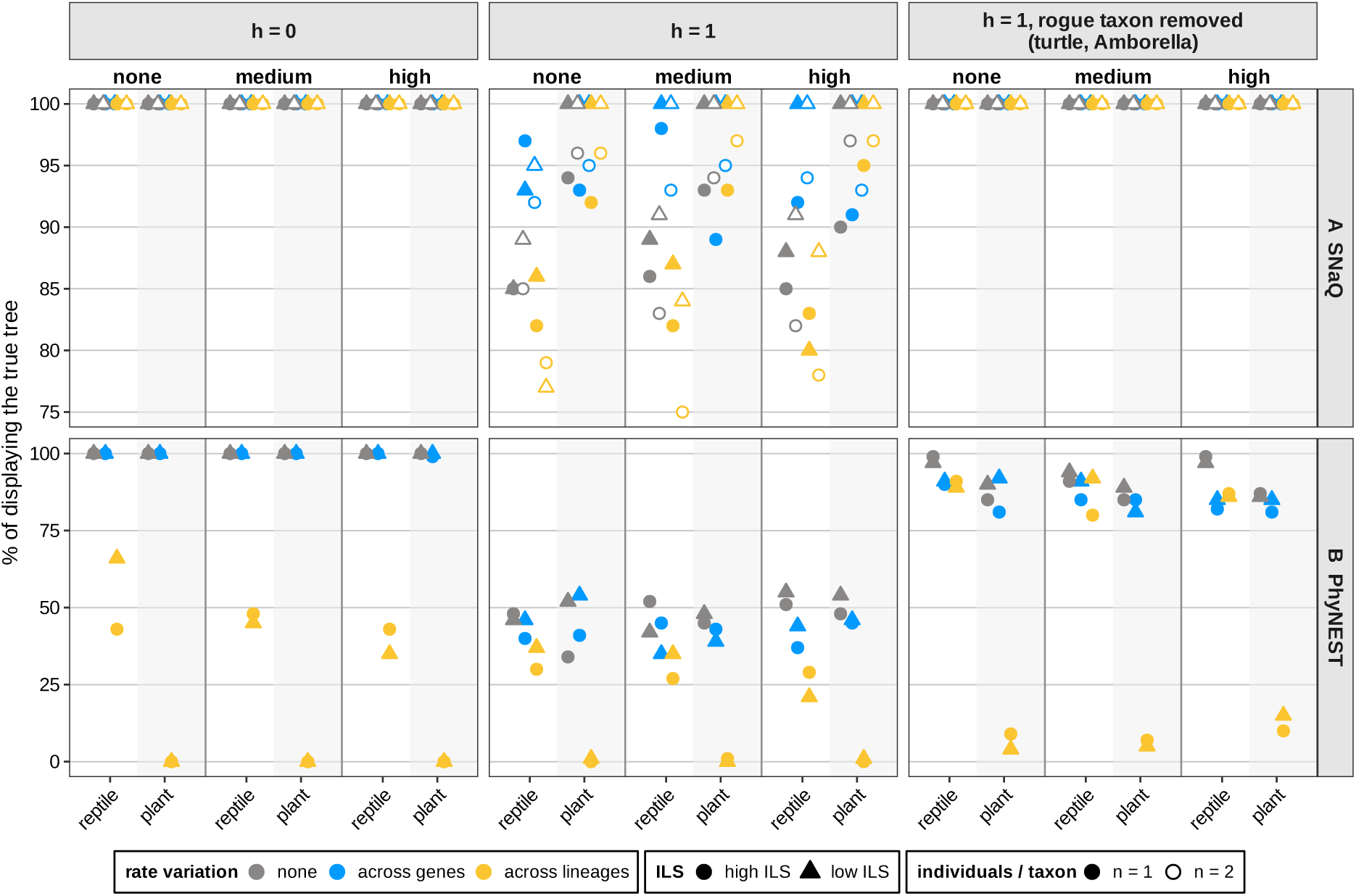
Probability that the true species topology is displayed in the inferred topology from (A) SNaQ and (B) PhyNEST (one individual per taxon). Note that the y-axis has different scales across panels. Columns, subpanels, colors, and shapes are as in Fig. E1.

## Supplementary Material F. Estimated proportion of gene flow when reticulation is allowed

**Figure F1.**
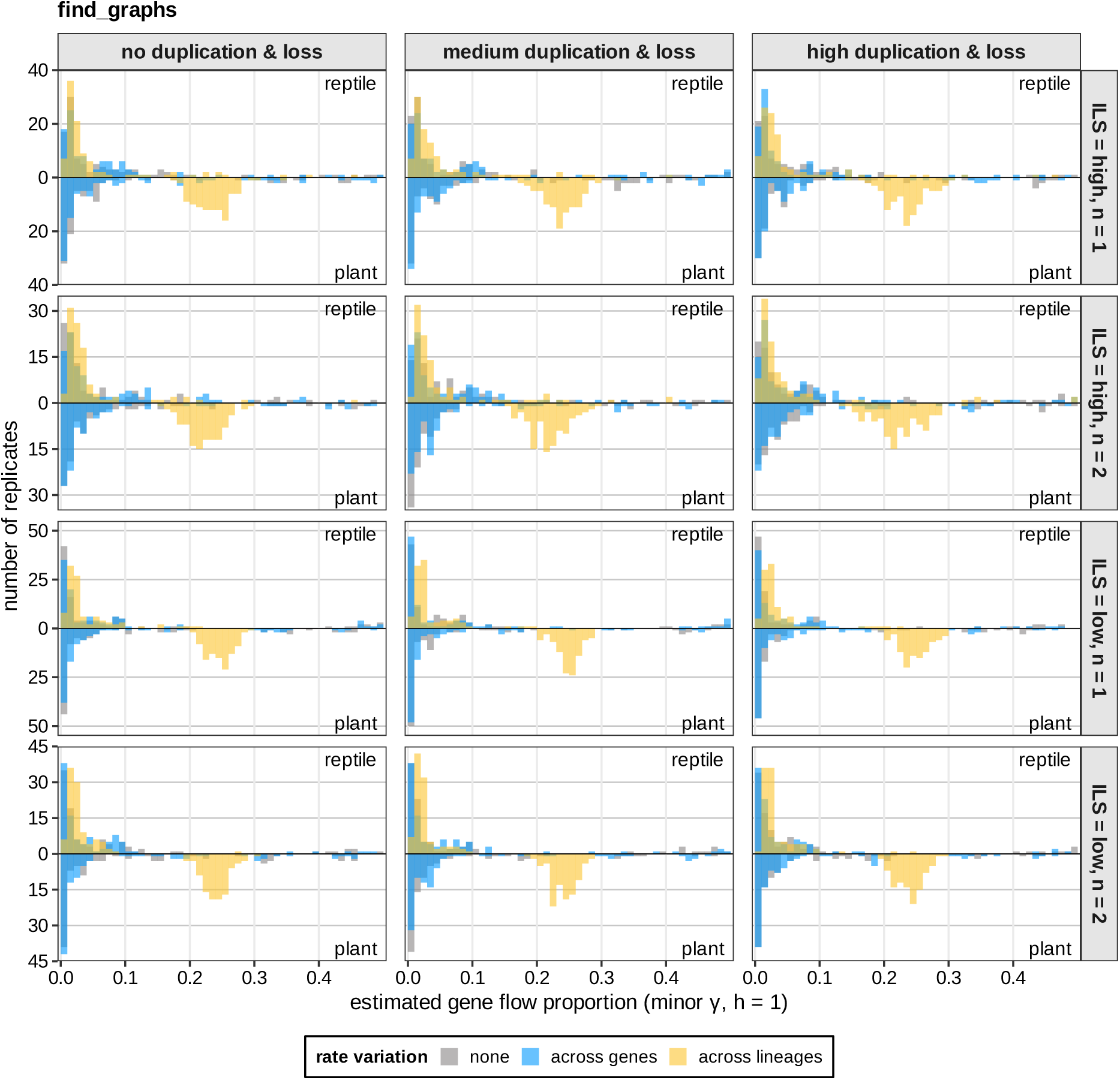
Estimated gene flow proportion (*γ*) inferred by find_graphs under *h* = 1. Columns correspond to duplication/loss levels (none, medium, high). Rows represent combinations of incomplete lineage sorting (ILS = high vs. low) and the number of sampled individuals per taxon (1 or 2). In each panel, the histogram of the reptile simulations is drawn upward and that of the plant simulations downward from the common horizontal zero line (counts of replicates, 100 per tree and setting). Histograms summarize estimates across simulation replicates under three substitution-rate models: rate variation across genes (blue), rate variation across lineages (yellow), and no rate variation (gray); the colors overlap.

**Figure F2.**
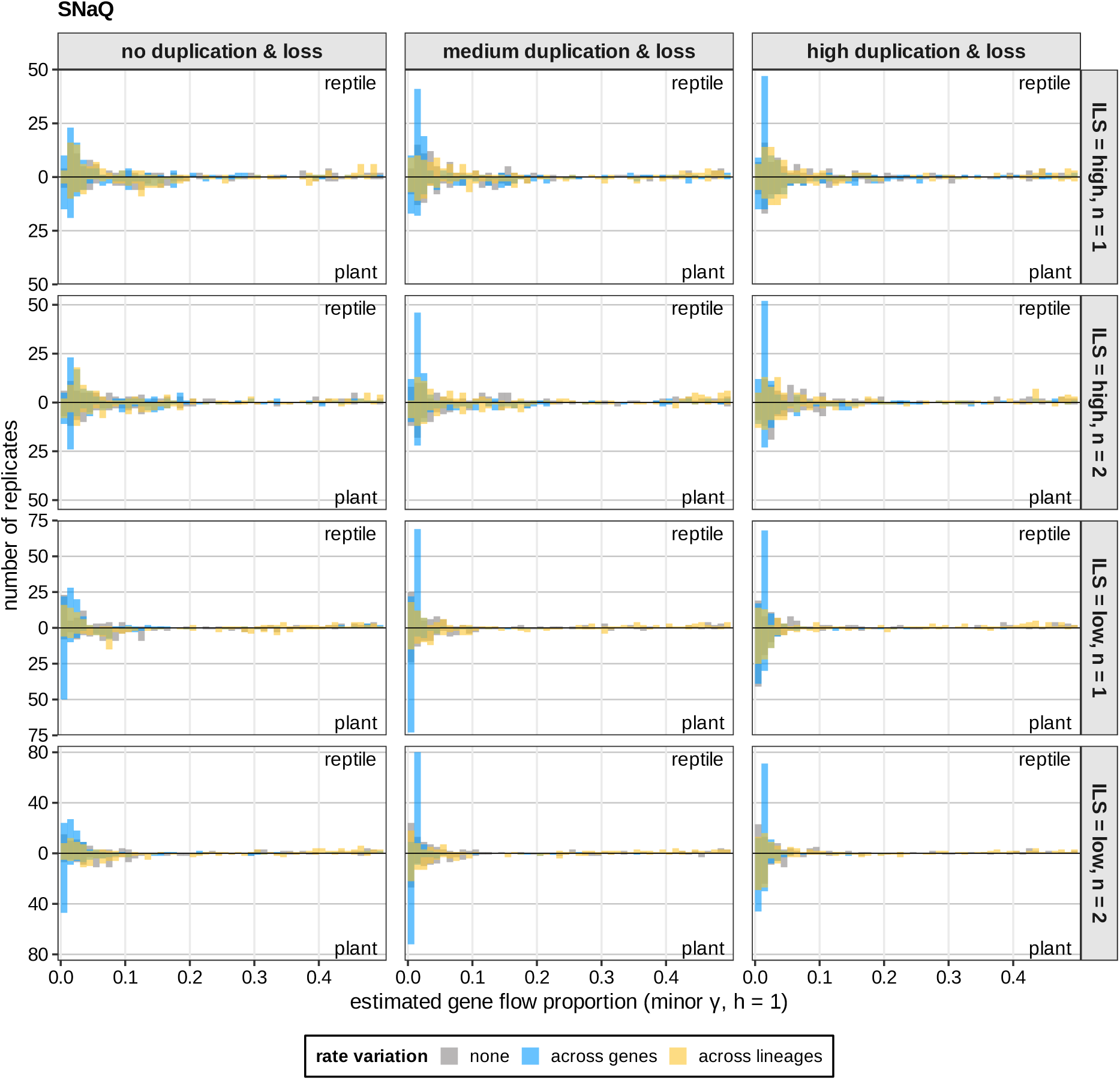
Estimated gene flow proportion (*γ*) of the *h* = 1 network inferred by SNaQ. Histograms of the estimates across the simulation replicates, the reptile tree drawn upward and the plant species tree downward from a shared horizontal zero line. Columns, rows and colors as in Fig. F1

**Figure F3.**
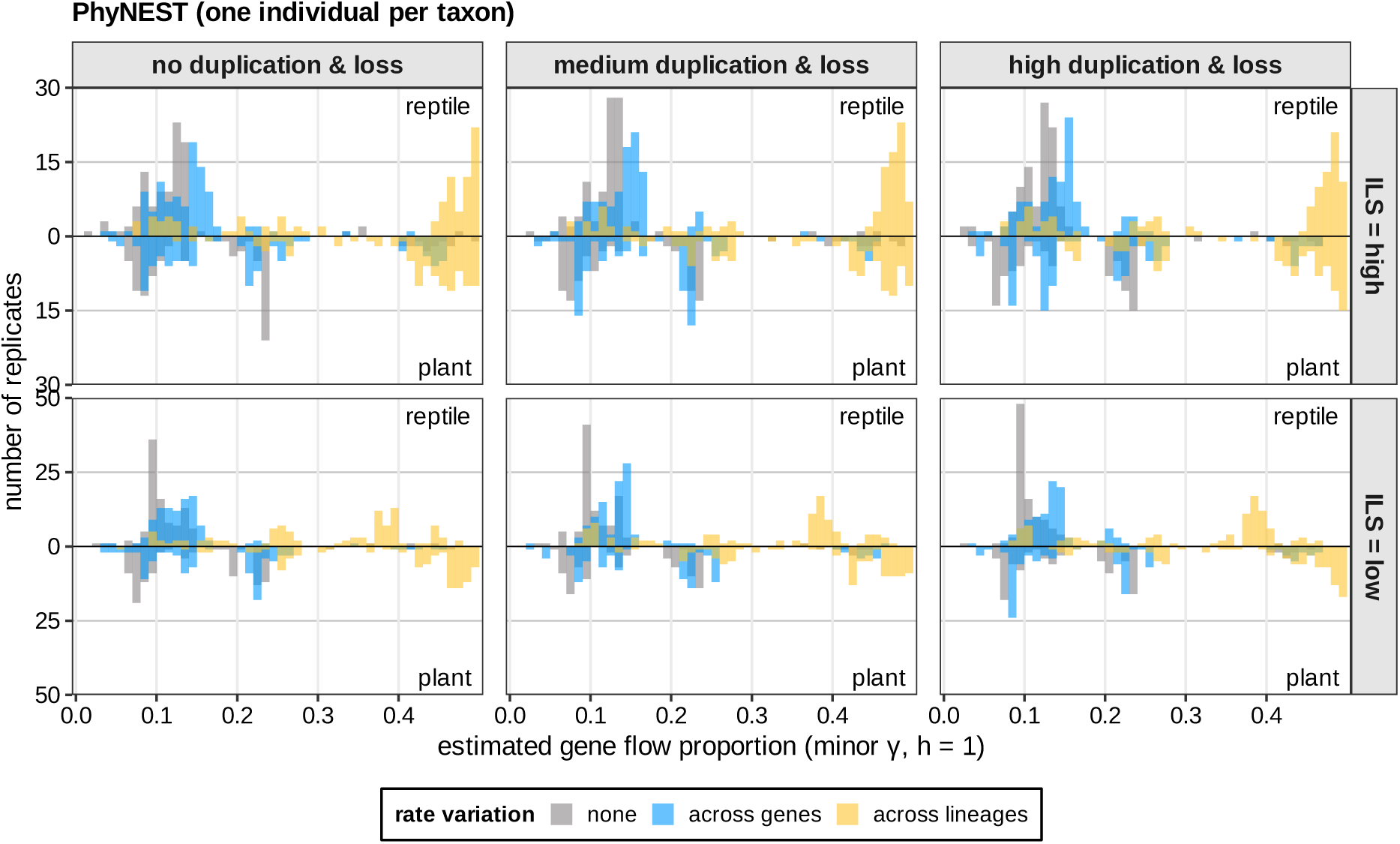
Estimated gene flow proportion (*γ*) of the *h* = 1 network inferred by PhyNEST across simulation scenarios (one individual per taxon). Histograms of the estimates across the simulation replicates, the reptile tree drawn upward and the plant species tree downward from a shared horizontal zero line, under three substitution-rate models: rate variation across genes (blue), across lineages (yellow), and none (gray). Columns correspond to the duplication/loss level (none, medium, high). Rows represent the level of incomplete lineage sorting (ILS = high vs. low).

**Table F1.**
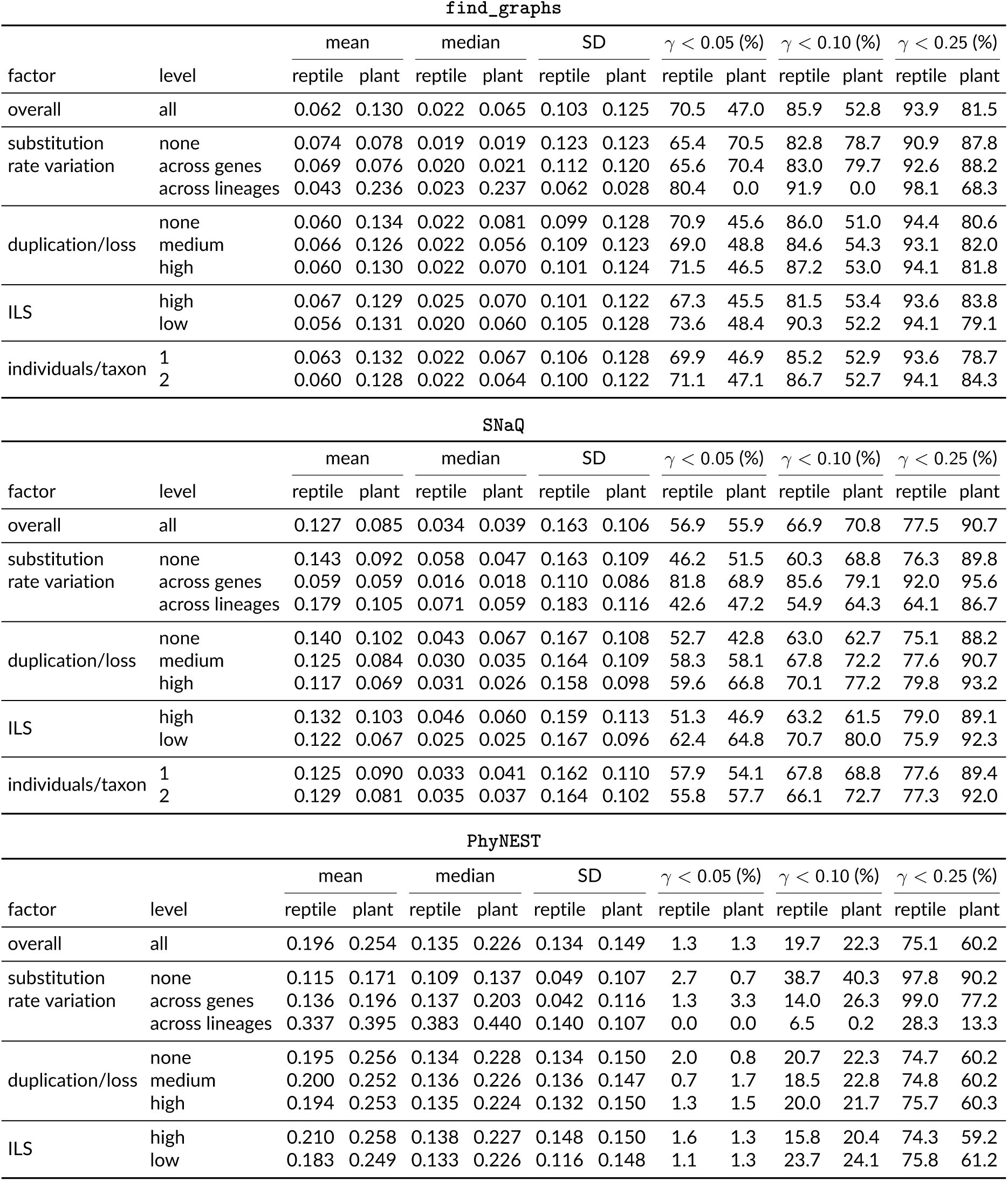
Summary statistics of estimated gene flow (minor *γ*) under *h* = 1 inferred by find_graphs (best-scoring graph), SNaQ and PhyNEST (one individual per taxon). For each factor level, the *γ* values from all replicates sharing that level are pooled across all other factor settings, and statistics are computed on the pooled values (100 replicates per setting; 36 settings per tree for find_graphs and SNaQ, 18 for PhyNEST). Percentages indicate the proportion of pooled *γ* values falling below each threshold.

| find_graphs |  |  |  |  |  |  |  |  |  |  |  |  |  |
|---|---|---|---|---|---|---|---|---|---|---|---|---|---|
|  |  | mean |  | median |  | SD |  | γ < 0.05 (%) |  | γ < 0.10 (%) |  | γ < 0.25 (%) |  |
| factor | level | reptile | plant | reptile | plant | reptile | plant | reptile | plant | reptile | plant | reptile | plant |
| overall | all | 0.062 | 0.130 | 0.022 | 0.065 | 0.103 | 0.125 | 70.5 | 47.0 | 85.9 | 52.8 | 93.9 | 81.5 |
| substitution rate variation | none | 0.074 | 0.078 | 0.019 | 0.019 | 0.123 | 0.123 | 65.4 | 70.5 | 82.8 | 78.7 | 90.9 | 87.8 |
|  | across genes | 0.069 | 0.076 | 0.020 | 0.021 | 0.112 | 0.120 | 65.6 | 70.4 | 83.0 | 79.7 | 92.6 | 88.2 |
|  | across lineages | 0.043 | 0.236 | 0.023 | 0.237 | 0.062 | 0.028 | 80.4 | 0.0 | 91.9 | 0.0 | 98.1 | 68.3 |
| duplication/loss | none | 0.060 | 0.134 | 0.022 | 0.081 | 0.099 | 0.128 | 70.9 | 45.6 | 86.0 | 51.0 | 94.4 | 80.6 |
|  | medium | 0.066 | 0.126 | 0.022 | 0.056 | 0.109 | 0.123 | 69.0 | 48.8 | 84.6 | 54.3 | 93.1 | 82.0 |
|  | high | 0.060 | 0.130 | 0.022 | 0.070 | 0.101 | 0.124 | 71.5 | 46.5 | 87.2 | 53.0 | 94.1 | 81.8 |
| ILS | high | 0.067 | 0.129 | 0.025 | 0.070 | 0.101 | 0.122 | 67.3 | 45.5 | 81.5 | 53.4 | 93.6 | 83.8 |
|  | low | 0.056 | 0.131 | 0.020 | 0.060 | 0.105 | 0.128 | 73.6 | 48.4 | 90.3 | 52.2 | 94.1 | 79.1 |
| individuals/taxon | 1 | 0.063 | 0.132 | 0.022 | 0.067 | 0.106 | 0.128 | 69.9 | 46.9 | 85.2 | 52.9 | 93.6 | 78.7 |
|  | 2 | 0.060 | 0.128 | 0.022 | 0.064 | 0.100 | 0.122 | 71.1 | 47.1 | 86.7 | 52.7 | 94.1 | 84.3 |
| SNaQ |  |  |  |  |  |  |  |  |  |  |  |  |  |
|  |  | mean |  | median |  | SD |  | γ < 0.05 (%) |  | γ < 0.10 (%) |  | γ < 0.25 (%) |  |
| factor | level | reptile | plant | reptile | plant | reptile | plant | reptile | plant | reptile | plant | reptile | plant |
| overall | all | 0.127 | 0.085 | 0.034 | 0.039 | 0.163 | 0.106 | 56.9 | 55.9 | 66.9 | 70.8 | 77.5 | 90.7 |
| substitution rate variation | none | 0.143 | 0.092 | 0.058 | 0.047 | 0.163 | 0.109 | 46.2 | 51.5 | 60.3 | 68.8 | 76.3 | 89.8 |
|  | across genes | 0.059 | 0.059 | 0.016 | 0.018 | 0.110 | 0.086 | 81.8 | 68.9 | 85.6 | 79.1 | 92.0 | 95.6 |
|  | across lineages | 0.179 | 0.105 | 0.071 | 0.059 | 0.183 | 0.116 | 42.6 | 47.2 | 54.9 | 64.3 | 64.1 | 86.7 |
| duplication/loss | none | 0.140 | 0.102 | 0.043 | 0.067 | 0.167 | 0.108 | 52.7 | 42.8 | 63.0 | 62.7 | 75.1 | 88.2 |
|  | medium | 0.125 | 0.084 | 0.030 | 0.035 | 0.164 | 0.109 | 58.3 | 58.1 | 67.8 | 72.2 | 77.6 | 90.7 |
|  | high | 0.117 | 0.069 | 0.031 | 0.026 | 0.158 | 0.098 | 59.6 | 66.8 | 70.1 | 77.2 | 79.8 | 93.2 |
| ILS | high | 0.132 | 0.103 | 0.046 | 0.060 | 0.159 | 0.113 | 51.3 | 46.9 | 63.2 | 61.5 | 79.0 | 89.1 |
|  | low | 0.122 | 0.067 | 0.025 | 0.025 | 0.167 | 0.096 | 62.4 | 64.8 | 70.7 | 80.0 | 75.9 | 92.3 |
| individuals/taxon | 1 | 0.125 | 0.090 | 0.033 | 0.041 | 0.162 | 0.110 | 57.9 | 54.1 | 67.8 | 68.8 | 77.6 | 89.4 |
|  | 2 | 0.129 | 0.081 | 0.035 | 0.037 | 0.164 | 0.102 | 55.8 | 57.7 | 66.1 | 72.7 | 77.3 | 92.0 |
| PhyNEST |  |  |  |  |  |  |  |  |  |  |  |  |  |
|  |  | mean |  | median |  | SD |  | γ < 0.05 (%) |  | γ < 0.10 (%) |  | γ < 0.25 (%) |  |
| factor | level | reptile | plant | reptile | plant | reptile | plant | reptile | plant | reptile | plant | reptile | plant |
| overall | all | 0.196 | 0.254 | 0.135 | 0.226 | 0.134 | 0.149 | 1.3 | 1.3 | 19.7 | 22.3 | 75.1 | 60.2 |
| substitution rate variation | none | 0.115 | 0.171 | 0.109 | 0.137 | 0.049 | 0.107 | 2.7 | 0.7 | 38.7 | 40.3 | 97.8 | 90.2 |
|  | across genes | 0.136 | 0.196 | 0.137 | 0.203 | 0.042 | 0.116 | 1.3 | 3.3 | 14.0 | 26.3 | 99.0 | 77.2 |
|  | across lineages | 0.337 | 0.395 | 0.383 | 0.440 | 0.140 | 0.107 | 0.0 | 0.0 | 6.5 | 0.2 | 28.3 | 13.3 |
| duplication/loss | none | 0.195 | 0.256 | 0.134 | 0.228 | 0.134 | 0.150 | 2.0 | 0.8 | 20.7 | 22.3 | 74.7 | 60.2 |
|  | medium | 0.200 | 0.252 | 0.136 | 0.226 | 0.136 | 0.147 | 0.7 | 1.7 | 18.5 | 22.8 | 74.8 | 60.2 |
|  | high | 0.194 | 0.253 | 0.135 | 0.224 | 0.132 | 0.150 | 1.3 | 1.5 | 20.0 | 21.7 | 75.7 | 60.3 |
| ILS | high | 0.210 | 0.258 | 0.138 | 0.227 | 0.148 | 0.150 | 1.6 | 1.3 | 15.8 | 20.4 | 74.3 | 59.2 |
|  | low | 0.183 | 0.249 | 0.133 | 0.226 | 0.116 | 0.148 | 1.1 | 1.3 | 23.7 | 24.1 | 75.8 | 61.2 |

